# An immune-stimulating antibody conjugate spatiotemporally targeting CD47 and TLR9 elicits macrophage-dependent tumor clearance and durable anti-cancer adaptive immunity

**DOI:** 10.64898/2026.07.22.739359

**Authors:** Byoungjae Kong, Seung Woo Chung, Daiheon Lee, Gijung Kwak, Jung Soo Suk

## Abstract

Cluster of differentiation 47 (CD47) blockade promotes tumor cell phagocytosis; however, transitioning this initial innate immune event into durable anti-cancer adaptive immunity and tumor control remains a critical challenge. To address this translational gap, we have engineered aCD47 – CpG, a novel immune-stimulating antibody conjugate (ISAC) coupling a CD47-blocking antibody (aCD47) to a Toll-like receptor 9 agonist, unmethylated CpG, to enable synchronized deployment of co-stimulatory signals during tumor engulfment. This ISAC, aCD47-CpG, but not the parent aCD47, reprograms macrophages toward an anti-tumor M1-like phenotype, enhances antigen cross-presentation, and promotes robust CD8^+^ T cell priming. We found that systemically administered aCD47-CpG drove macrophage-dependent tumor suppression in a human lymphoma xenograft model. In parallel, the treatment in immunocompetent syngeneic models of lymphoma and highly immunosuppressive triple-negative breast cancer resulted in profound tumor regression, metastasis inhibition, and durable anti-cancer immune memory. These effects were accompanied by remodeling of the immunosuppressive tumor microenvironment toward an immune-permissive state, characterized by M2-to-M1 macrophage repolarization, intratumoral infiltration of cytotoxic and memory T cells, and depletion of regulatory T cells. This spatiotemporally coordinated immunotherapy platform offers a promising strategy to bridge innate and adaptive immunity, thereby advancing the translational potential of CD47-targeted cancer therapies.

## Main

Cluster of differentiation 47 (CD47) is a ubiquitous macrophage checkpoint molecule upregulated across diverse hematologic and solid tumors to evade phagocytosis^1–3^. Although CD47-blocking therapies, such as anti-CD47 antibodies (aCD47), have shown potent preclinical efficacy and tolerable safety profiles in humans^4–9^, they have yielded inconsistent clinical responses and often failed to drive durable tumor regression^5, 6, 8, 9^. This clinical limitation underscores that promoting phagocytosis alone is insufficient and that durable tumor control requires concurrent engagement of cancer-specific adaptive immunity^10, 11^.

Achieving this therapeutic goal hinges upon linking initial tumor cell engulfment with cross-priming of tumor-specific CD8^+^ T cells^12, 13^. CD47 blockade has shown ability to induce cross-priming in highly immunogenic preclinical models^12, 14, 15^. However, eliciting T cell activation against poorly immunogenic endogenous tumor antigens remains a formidable clinical challenge, because tumor cell engulfment typically results in immunologically silent tumor clearance^16^. This inherent tolerance is further compounded by the highly immunosuppressive tumor microenvironment (TME), which is primarily dominated by pro-tumorigenic M2-like macrophages and regulatory T cells (Tregs) that collectively restrain anti-tumor immunity^17–19^. Incorporation of potent co-stimulatory stimuli capable of breaking myeloid tolerance, reverting immunosuppression, and instructing macrophages for optimal cross-presentation^20^ represents a compelling strategy to overcome these translational gaps.

Systemic co-administration of aCD47 and co-stimulatory agents is a logical clinical strategy, but this approach is fundamentally limited by a critical spatiotemporal barrier; uncoordinated delivery of individual components cannot ensure sequential bundling of tumor cell engulfment and co-stimulatory signal activation by identical macrophages. We hypothesized that this spatiotemporal synchronization can be achieved by simultaneous delivery of aCD47 and co-stimulatory agents as a chemically coupled single entity. This spatial requirement makes Toll-like receptor 9 (TLR9) an ideal target, because it is strictly localized within intracellular endosomes and phagolysosomes^21^ where engulfed tumor antigens are processed. To exploit this target, we have selected a TLR9 agonist, synthetic unmethylated CpG oligodeoxynucleotide (CpG), as a potent co-stimulatory payload. CpG drives Th1-polarized immunity and orchestrates robust CD8^+^ T cell cross-priming^22^ while its clinical safety has been established in clinical studies^22, 23^.

Accordingly, we have engineered aCD47-CpG, a novel immune-stimulating antibody conjugate (ISAC) comprising aCD47 covalently linked to CpG. This platform couples aCD47-mediated tumor cell engulfment with phagolysosomal CpG delivery to ensure that tumor-engulfing macrophages receive intracellular co-stimulatory signals essential for efficient antigen cross-presentation. Here, we present rational design and mechanistic validation of this ISAC using complementary *in vitro* and *in vivo* models. In addition to *in vitro* co-culture models, we utilized human xenograft and immunocompetent syngeneic tumor models to assess innate-only macrophage-dependent tumor control and orchestrated anti-cancer innate and adaptive immune responses, respectively.

## Results

### aCD47-CpG retains binding affinity to individual targets while promoting NF-κB activity *in vitro*

To identify the most effective aCD47-CpG conjugate, we engineered four candidates with varying drug-antibody ratios (DARs) ranging from 1 to 4 (**Fig. 1a**). This was achieved through a two-step reaction using non-cleavable heterobifunctional crosslinker containing N-hydroxysuccinimide (NHS)– ester and maleimide reactive groups, which react with the primary amines of mouse aCD47 and sulfhydryl groups added to the 3’-end of CpG, respectively (**Fig. 1b**). CpG molecules were conjugated via their 3’-end to preserve immunostimulatory activity, as 5’-end modification has been reported to reduce the potency^24^. To examine the conjugation pattern, we visualized the products via SDS-PAGE, utilizing Coomassie blue staining for aCD47 and SYBR Gold staining for CpG. We observed DAR-dependent molecular weight increments of the target protein bands, reflecting distinct CpG conjugation to aCD47 (**Fig. 1c**). This finding was further validated via size exclusion chromatography, where conjugates with higher DARs eluted earlier, indicating increased molecular size (**Fig. 1d**). Given that chemical modification of aCD47 by CpG addition may influence CD47 binding affinity, we evaluated the binding of aCD47-CpG to mouse CD47 (mCD47) and TLR9 (mTLR9) via ELISA. The analysis revealed that the binding affinity of the conjugates to mCD47 was well retained at nanomolar levels, exhibiting only a marginal, DAR-dependent increase in the K_D_ value compared to the aCD47 control (**Supplementary Table 1**), likely attributed to steric hindrance. The binding affinity of aCD47-CpGs to mTLR9 was also retained with an expected DAR dependency (**Fig. 1e**). Next, we explored the ability of aCD47-CpG to induce the activation of NF-κB in macrophages, a key regulator that controls innate and adaptive immune responses at multiple levels in cancer-related inflammation^25^. To do this, we leveraged RAW-Blue cells, which express a secreted embryonic alkaline phosphatase (SEAP) reporter gene driven by NF-κB and AP-1 activation upon TLR9 stimulation. We measured SEAP activity in the supernatants after treating RAW-Blue cells with aCD47-CpG candidates in the presence or absence of A20 mouse lymphoma cells. All tested aCD47-CpG conjugates significantly induced NF-κB activity in mono-cultures of mouse macrophages, confirming the preserved biological function of CpG within the conjugate (**Fig. 1f**). Notably, NF-κB activity was further elevated in co-cultures with A20 cells, suggesting that increased uptake of aCD47-CpG by RAW-Blue cells was mediated by CD47-overexpressing A20 cells (**Fig. 1f** and **Supplementary Fig. 1**). However, NF-κB signaling plateaued beyond a DAR of 3, with no significant difference observed compared to a DAR of 4. We therefore selected the conjugate with a DAR of 3 for further evaluation, as it represents the optimal stoichiometry that maximizes TLR9 stimulation while retaining CD47 binding. To establish the clinical translatability of our platform, we applied this optimized conjugation strategy to generate a human-version aCD47-CpG comprising human aCD47 and CpG, which retained robust target engagements and TLR9-stimulating ability (**Extended Data Fig. 1**), similar to our observations with mouse aCD47-CpG (**Fig. 1**).

**Fig. 1.**
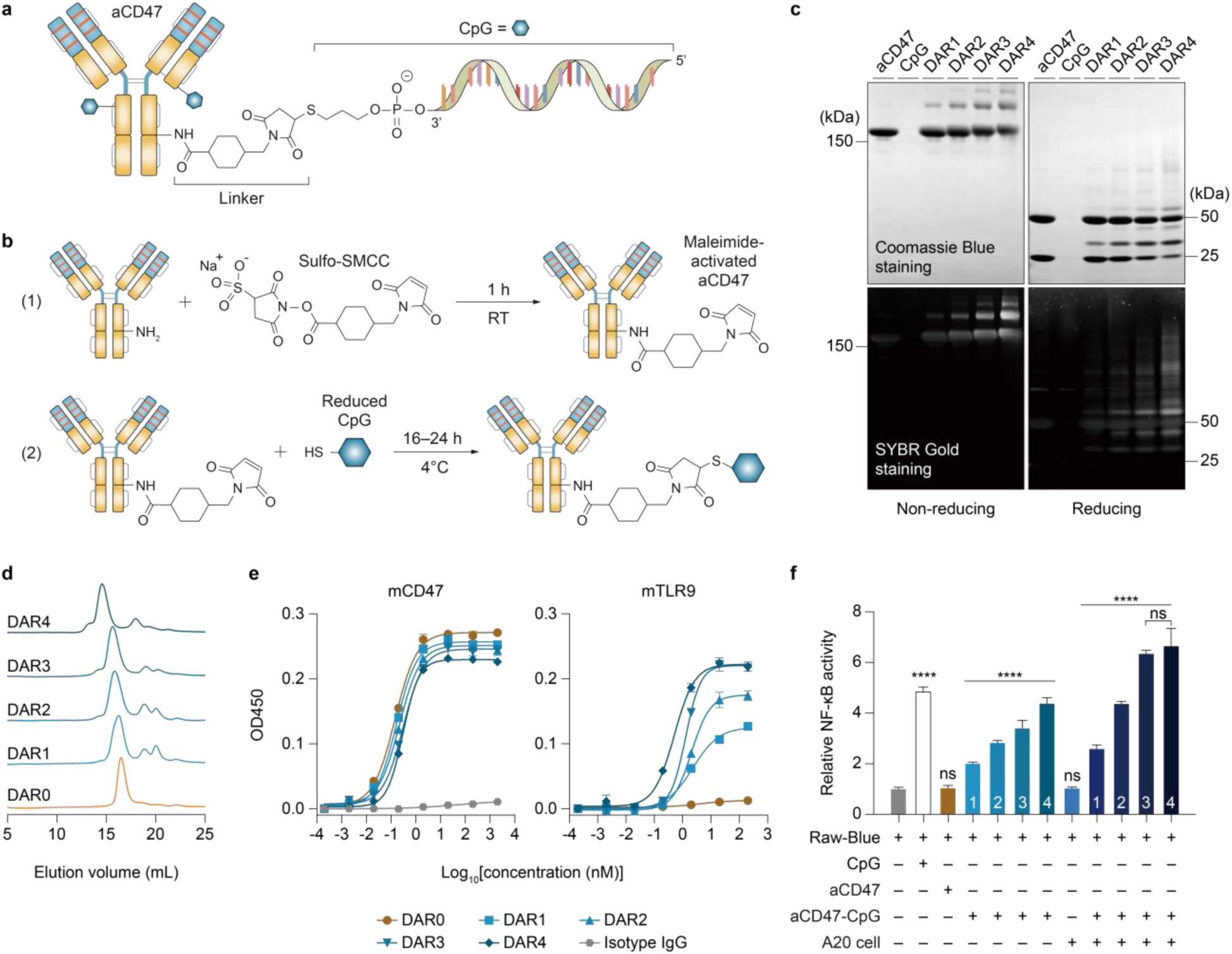
| Mouse aCD47-CpG retains binding affinity to CD47 and TLR9 while promoting NF-κB activity *in vitro*. **a**, Schematic illustration of aCD47-CpG conjugate. **b**, Graphical depiction of two-step conjugation reaction for aCD47-CpG synthesis. RT, room temperature. **c**, SDS-PAGE analysis for conjugation patterns of aCD47-CpGs engineered at drug-antibody ratios (DARs) of 1–4 under non-reducing (left) and reducing (right) conditions. Upper and lower two gels show Coomassie Blue and SYBR Gold staining for the visualization of antibodies and CpG, respectively. **d**, Size exclusion chromatography analysis of aCD47-CpG candidates. **e**, Binding curves of aCD47-CpG candidates to recombinant mouse CD47 (mCD47, left) and TLR9 (mTLR9, right). **f**, NF-κB activities induced by aCD47-CpG candidates. Wherever relevant, Raw-Blue cells were stimulated with CpG or aCD47-CpG at a CpG concentration of 1 μg/mL in the presence or absence of mouse lymphoma A20 cells. Individual DAR values are indicated inside the respective bars. Data shown are mean ± s.d. of three experimental replicates. ns: non-significant, \*\*\*\**p* < 0.0001 (one-way ANOVA with Tukey’s multiple comparisons test).

### aCD47-CpG enhances cancer cell phagocytosis by macrophages *in vitro*

CD47 blockade strategy has been most widely explored against hematologic tumors in clinical studies, and its tolerability has been established among patients with lymphoma^26^. We thus investigated the ability of the optimized aCD47-CpG conjugate to promote macrophage-mediated phagocytosis of non-Hodgkin lymphoma (NHL) cells. To do so, we conducted a conventional *in vitro* phagocytosis assay by co-culturing GFP-expressing mouse lymphoma A20 (A20-GFP) cells with syngeneic BALB/c bone marrow-derived macrophages (BMDMs) in the presence of aCD47-CpG for 2 h at 37°C. To efficiently target mCD47 on A20 cells, aCD47-CpG was generated using clone MIAP410, a well-characterized anti-mCD47 antibody known to functionally block mouse CD47-SIRPα interactions^3^. In parallel, a human aCD47-CpG was prepared using clone B6H12 to target human CD47, thereby enabling phagocytosis of GFP-expressing human Raji lymphoma (Raji-GFP) cells. Unconjugated mouse aCD47 induced modest but significant phagocytosis of A20 cells (**Fig. 2a**) whereas mouse aCD47-CpG did not, presumably reflecting its attenuated CD47-binding affinity demonstrated by ELISA (**Fig. 1e**). Conversely, both human aCD47 and aCD47-CpG exhibited robust phagocytosis of Raji-GFP cells (**Fig. 2a**), consistent with previous reports^2^. These distinct phagocytic patterns were further confirmed via microscopy-based visualization (**Fig. 2b**). To gain deeper insight into the sustained phagocytic potential of aCD47-CpG, we conducted a luminescence-based phagocytosis assay, with an extended incubation time beyond 2 h. Unlike conventional flow-based phagocytosis assays that quantify engulfing macrophages, this method measures the surviving luciferase-expressing cancer cells to provide enhanced sensitivity and precision^27^. Remarkably, following this prolonged incubation, mouse aCD47-CpG significantly enhanced A20 phagocytosis compared to aCD47 alone (**Fig. 2c**), and this effect was independent of any direct impact on luciferase activity (**Supplementary Fig. 2**). To dissect the temporal dynamics of this process, we conducted a kinetic analysis of BMDM-mediated phagocytosis. During the early stages of incubation, aCD47 significantly induced phagocytosis of A20 cells, whereas aCD47-CpG was unable to do so, corroborating the observation in the flow-based phagocytosis assay (**Fig. 2a**). However, beginning at 8-h post-incubation, phagocytosis mediated by aCD47-CpG surpassed that induced by aCD47 (**Fig. 2d**), underscoring the critical role of CpG in promoting sustained phagocytic activity^28^. The prolonged exposure to aCD47-CpG markedly upregulated co-stimulatory molecules, including CD80, CD86, and CD40, and significantly increased the expression of iNOS, a hallmark of M1-like macrophage polarization^29^, in BMDMs (**Fig. 2e**). Moreover, aCD47-CpG-stimulated BMDMs strongly released pro-inflammatory cytokines, including TNFα, IL-6, and IL-12 (**Fig. 2f**), which are key mediators bridging innate and adaptive immune responses^30^. The human version of aCD47-CpG similarly elicited robust phagocytosis of Raji cells in prolonged incubation, as visually confirmed by fluorescence microscopy (**Extended Data Fig. 2a,b**). Macrophages stimulated with this formulation exhibited enhanced expression of the co-stimulatory molecule CD40 and robust secretion of pro-inflammatory cytokines, paralleling the responses observed in the mouse co-culture model (**Extended Data Fig. 2c,d**). Collectively, these findings demonstrate that aCD47-CpG not only improves NHL cell phagocytosis but also extensively reprograms macrophages to augment anti-tumor immunity.

**Fig. 2.**
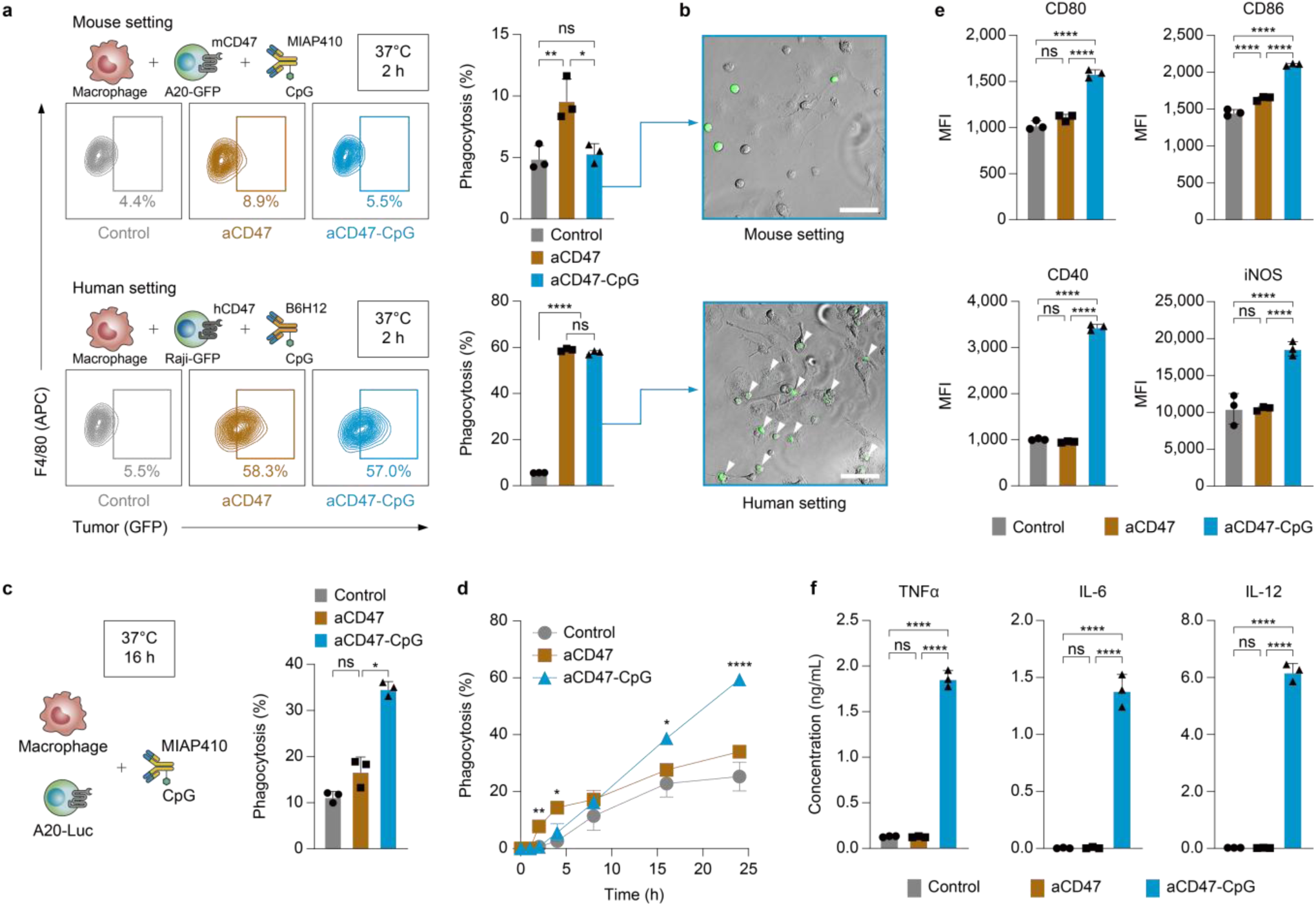
| aCD47-CpG promotes phagocytosis and anti-tumor immune response of macrophages against NHL cells *in vitro*. **a–f**, Bone marrow-derived macrophages (BMDMs) were co-cultured with lymphoma cell lines at a 2:1 ratio in the presence of aCD47 or aCD47-CpG. **a**, Representative flow cytometry plots elucidating the phagocytosis of GFP-expressing mouse A20 cells (A20-GFP) or human Raji cells (Raji-GFP) by macrophages for 2 h at 37°C. MIAP410 and B6H12 antibody clones specific to mouse and human CD47, respectively, were utilized. Macrophages were stained with APC-labeled anti-F4/80 antibodies. **b**, Representative phase contrast images of aCD47-CpG-treated co-cultures as in **a**. Scale bars = 50 µm. Phagocytosed tumor cells were indicated by white arrows. **c**, Phagocytosis for 16 h at 37°C determined by luminescence-based long-term macrophage killing (LB-LTMK) assay. **d**, Kinetics of aCD47– or aCD47-CpG-mediated phagocytosis determined by LB-LTMK assay. **e**, Expression of co-stimulatory molecules (CD80, CD86, and CD40) and iNOS from BMDMs co-cultured with A20 cells after treatment with aCD47 or aCD47-CpG. Cells were fixed and permeabilized prior to flow cytometry analysis for the quantification of intracellular iNOS level. **f**, Pro-inflammatory cytokines released from BMDMs by aCD47-CpG stimulation. The levels of TNFα, IL-6, or IL-12 secreted to cell culture supernatants were measured by ELISA after 24-h incubation. Data shown are mean ± s.d. of three experimental replicates. ns: non-significant, \**p* < 0.05, \*\**p* < 0.01, \*\*\*\**p* < 0.0001 (one-way ANOVA with Tukey’s multiple comparisons test).

### aCD47-CpG elevates antigen cross-presentation and T cell priming *in vitro*

We next examined whether aCD47-CpG-activated BMDMs could effectively cross-present tumor antigens to CD8^+^ T cells. To ensure MHC class I (MHC-I) compatibility with the OT-I T cell system, we utilized BMDMs derived from C57BL/6 mice and co-cultured them with ovalbumin-expressing E.G7 (E.G7-OVA), a syngeneic mouse lymphoma cell line. Following overnight incubation with mouse aCD47-CpG, antigen cross-presentation was evaluated by detecting MHC-I-bound SIINFEKL peptides on the BMDMs (**Fig. 3a,b**). aCD47-CpG significantly augmented antigen cross-presentation, whereas aCD47 failed to elicit this effect (**Fig. 3b**). Likewise, the human aCD47-CpG markedly promoted antigen cross-presentation in BMDMs co-incubated with Raji cells expressing ovalbumin (Raji-OVA) (**Extended Data Fig. 2e**). To determine whether this enhanced cross-presentation translates to efficient T cell priming, we measured OVA-specific CD8^+^ T cell proliferation using a carboxyfluorescein succinimidyl ester (CFSE)-dilution assay (**Fig. 3a**). We co-cultured BMDMs with E.G7-OVA cells in the presence of control, aCD47, or aCD47-CpG, followed by the addition of naïve CD8^+^ T cells isolated from OT-I transgenic mice. Treatment with aCD47-CpG significantly enhanced the proliferation of OT-I T cells compared to aCD47, as quantified by CFSE dilution on day 3 (**Fig. 3c**). Importantly, aCD47-CpG-stimulated BMDMs elicited robust IFN-γ secretion from co-cultured CD8^+^ T cells, whereas aCD47 yielded negligible IFN-γ production (**Fig. 3d**). Collectively, these findings demonstrate that aCD47-CpG not only enhances tumor cell phagocytosis but also promotes antigen-specific CD8^+^ T cell priming, effectively bridging innate engulfment with robust adaptive anti-tumor immunity.

**Fig. 3.**
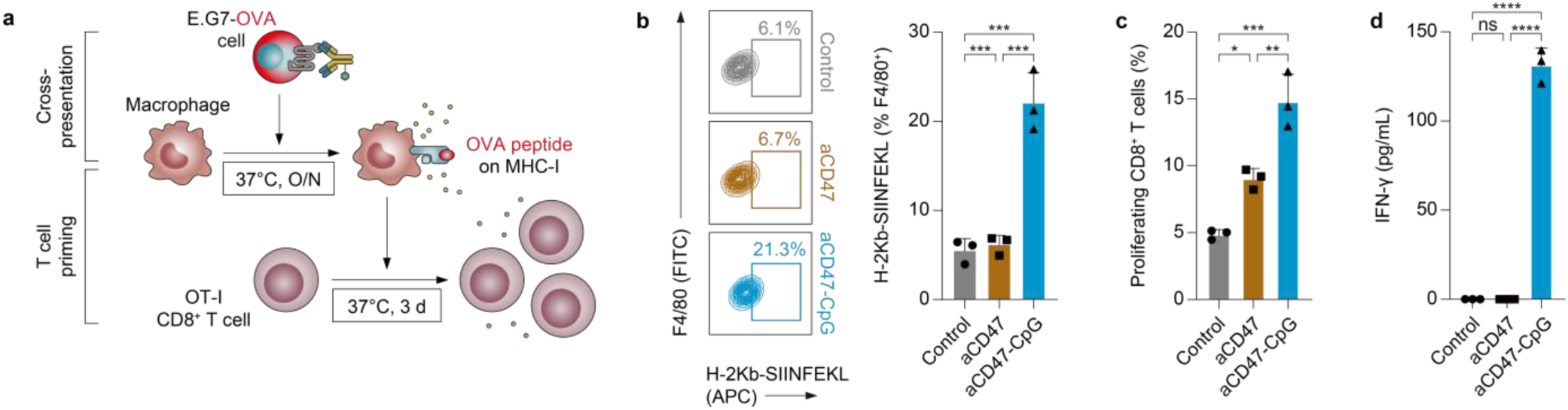
| aCD47-CpG elevates antigen cross-presentation and T cell priming *in vitro*. **a**, Schematics showing the experimental design for assessing cross-presentation and T cell priming. **b**, Cross-presentation of chicken ovalbumin-derived SIINFEKL peptide loaded onto H-2Kb (MHC-I) of macrophages. BMDMs were co-cultured with E.G7 cells expressing ovalbumin (E.G7-OVA) in the presence of aCD47 or aCD47-CpG overnight (O/N) at 37°C. Antigen cross-presentation was determined using flow cytometry by detecting MHC-I-bound SIINFEKL peptide on total F4/80^+^ macrophages. **c**, CD8^+^ T cell priming induced by macrophages after phagocytosis of E.G7-OVA cells mediated by aCD47-CpG. CD8^+^ T cells isolated from OT-I transgenic mice and labeled with CFSE (0.5 μM) were added to the co-culture of BMDM and E.G7-OVA in the presence of aCD47-CpG. Analysis was performed by flow cytometry on day 3. **d**, IFN-γ secreted by OT-I CD8^+^ T cells to co-culture supernatants in **c**. Data shown are mean ± s.d. of three experimental replicates. ns: non-significant, \**p* < 0.05, \*\**p* < 0.01, \*\*\**p* < 0.001, \*\*\*\**p* < 0.0001 (one-way ANOVA with Tukey’s multiple comparisons test).

### Systemically administered aCD47-CpG shows tolerability without incurring significant toxicity *in vivo*

To determine the maximum tolerated dose (MTD) for subsequent efficacy studies, we conducted a dose escalation study in healthy BALB/c mice. Given the potential hematological toxicities of CD47 blockade, we evaluated cohorts with and without a clinically validated low-dose 1-mg/kg priming strategy^26^, followed by a maintenance dose of 5, 10, or 20 mg/kg (**Extended Data Fig. 3a**). Over the 3-week observation period, survival was 100% across all cohorts (**Extended Data Fig. 3b**). While PBS-treated control mice exhibited steady weight gain, all aCD47-CpG-treated cohorts displayed a transient, fully reversible delay in weight gain during the first 5 days, irrespective of the priming (**Extended Data Fig. 3c**). Hematological tracking in unprimed cohorts revealed a transient reduction in RBC counts within 2 hours post-injection across all groups, including the PBS control (**Extended Data Fig. 3d**). This uniform initial drop suggests a procedure-related physiological artifact, such as transient hemodilution or handling stress^31^, rather than an ISAC-specific toxicity. Importantly, RBC levels in all cohorts completely recovered to normal baselines by day 7 (**Extended Data Fig. 3e**). Although generally well-tolerated, mice that received the highest dose (20 mg/kg) exhibited clinical signs of distress (*e.g.*, hunched posture, reduced mobility). Their clinical scores exceeded 2, the predetermined threshold for dose limitation^32^. Consequently, we established 10 mg/kg as the MTD, as it demonstrated an optimal balance between systemic safety and tolerability without requiring a low-dose priming step.

### Systemically administered aCD47-CpG preferentially accumulates in tumors *in vivo*

To evaluate *in vivo* biodistribution, we established a syngeneic NHL model by subcutaneously implanting A20 cells. Once tumors reached ∼100–200 mm³, mice received an intravenous injection of PBS, Cy5-labeled Isotype-CpG, or Cy5-labeled aCD47-CpG (10 mg/kg) (**Extended Data Fig. 4a** and **Supplementary Fig. 3a**). At 24-h post-dosing, *ex vivo* imaging revealed that aCD47-CpG exhibited expected off-tumor accumulation in the lung and liver, consistent with ubiquitous CD47 antigen sinks^33^ and hepatic clearance, but its accumulation within the tumor vastly exceeded that in any other major organs (**Extended Data Fig. 4b,c** and **Supplementary Fig. 3b**). Furthermore, aCD47-CpG demonstrated significantly greater tumor retention compared to the isotype-CpG control (**Extended Data Fig. 4d,e**). Collectively, these findings confirm that specific CD47 engagement drives robust, tumor-targeted delivery despite the presence of systemic antigen sinks.

### Systemically administered aCD47-CpG elicits macrophage-dependent anti-tumor effects in a human xenograft model of NHL *in vivo*

To establish the therapeutic and clinical relevance of aCD47-CpG in an innate immune context, we prepared a human NHL xenograft model by subcutaneously implanting human Raji cells into BALB/c nude mice. Once tumors reached palpable sizes, mice were systemically treated with PBS, aCD47, or aCD47-CpG (10 mg/kg) every three days up to four doses (**Fig. 4a**). To mechanistically validate the macrophage dependency of our therapy, an additional cohort received aCD47-CpG combined with macrophage-depleting clodronate liposomes. Consistent with our *in vitro* cross-species findings, systemic administration of human aCD47-CpG resulted in profound tumor regression and a significant reduction in terminal tumor weights compared to aCD47 and PBS controls (**Fig. 4b,c**). This therapeutic efficacy was completely abolished in the macrophage-depleted cohort (**Fig. 4b,c**), indicating that macrophage-mediated innate immune activation was an indispensable primary driver of tumor clearance in this model. To further delineate immune modulation within the TME, we performed flow cytometry analysis using the harvested tumors. The overall frequency of intratumoral macrophages was comparable among all treatment groups without macrophage depletion while clodronate treatment markedly reduced it (**Fig. 4d**). Notably, treatment with aCD47-CpG, but not aCD47, rewired the macrophage population toward an M1-like phenotype, as evidenced by a marked increase in the M1-to-M2 ratio compared to both the PBS and aCD47 control groups (**Fig. 4e–g**), underscoring the ability of aCD47-CpG to reverse the immunosuppressive TME.

**Fig. 4.**
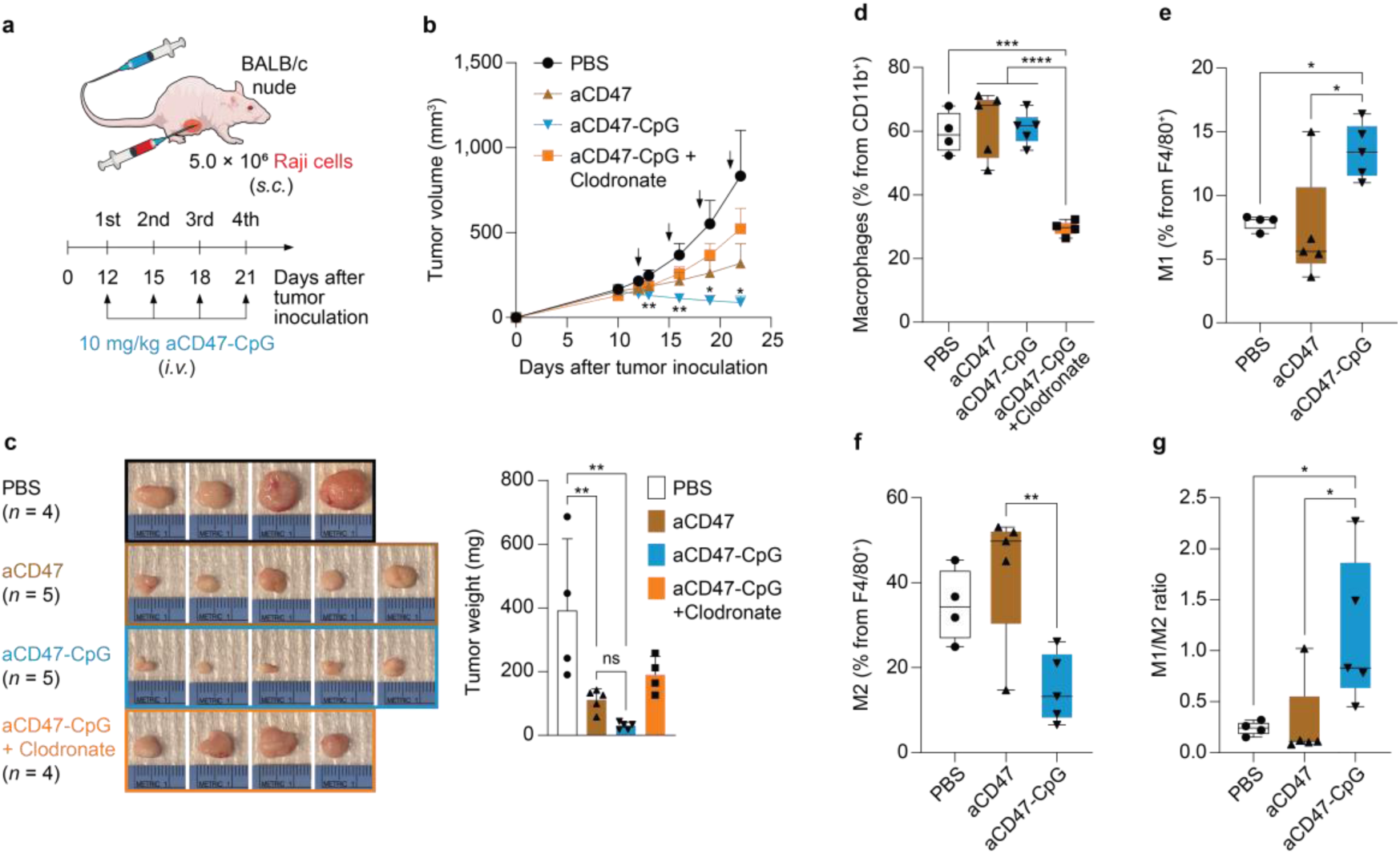
| Systemically administered human aCD47-CpG elicits macrophage-dependent anti-tumor effects in a human NHL xenograft model *in vivo*. **a**, Schematic diagram of tumor inoculation and treatment for a human lymphoma xenograft model. BALB/c nude mice were subcutaneously (*s.c.*) implanted with Raji cells. aCD47-CpG was given via intravenous administration (*i.v.*) at 10 mg/kg. For comparison and mechanistic validation, animals in different groups received PBS (control, *n* = 4), aCD47 (*n* = 5, 10 mg/kg), or aCD47-CpG with (*n* = 4, 10 mg/kg) or without (*n* = 5, 10 mg/kg) treatment with macrophage-depleting clodronate liposomes. **b**, Tumor growth of mice treated in **a**. Tumor progression was monitored by tumor volume measurement using a caliper. **c**, Images (left) and tumor weights (right) of excised Raji tumors at the endpoint. **d–g**, Flow cytometry analysis of macrophages in TME. The analysis highlights macrophage repolarization (**e–g**) within the total intratumoral CD11b^+^ cell population. Data shown are mean ± s.d. \**p* < 0.05, \*\**p* < 0.01, \*\*\**p* < 0.001, \*\*\*\**p* < 0.0001 (one-way ANOVA with Tukey’s multiple comparisons test).

### Systemically administered aCD47-CpG demonstrates robust anti-tumor therapeutic efficacy and immunological memory in a systemic syngeneic mouse model of NHL

Having established the therapeutic efficacy of aCD47-CpG against human tumors in an immunodeficient xenograft setting, we next sought to investigate its anti-tumor effects in an immunocompetent host with integral adaptive immune elements. We established a systemic syngeneic NHL mouse model and conducted efficacy studies at the predetermined MTD of 10 mg/kg. First, we intravenously injected luciferase-expressing A20 (A20-Luc) cells into BALB/c mice. Cyclophosphamide was systemically administered one day prior to tumor inoculation to facilitate efficient tumor engraftment while preserving lymphoreplete conditions, allowing for studies in a fully immunocompetent mouse model without inducing generalized lymphodepletion^34^. When systemic lymphoma was confirmed by live-animal bioluminescence imaging (BLI) (10 days later), mice were randomly assigned to three groups and systemically treated with PBS, aCD47, or aCD47-CpG (10 mg/kg) using the same dosing schedule (every third day up to four doses) (**Fig. 5a**). Relative to the PBS and aCD47 control groups, aCD47-CpG treatment markedly reduced the tumor burden (**Fig. 5b,c**) and significantly extended survival (**Fig. 5d**). To assess the establishment of long-term anti-cancer immunity, five surviving mice from the aCD47-CpG-treated group were rechallenged with A20 cells on day 106 at an identical cell count as the initial inoculation. Remarkably, these previously inoculated and treated mice were completely resistant to A20-cell rechallenge unlike the age-matched *de novo* inoculated control mice that succumb to death (**Fig. 5e**), demonstrating that aCD47-CpG elicited durable systemic anti-tumor immune memory to prevent tumor relapse. Collectively, these findings highlight aCD47-CpG as a highly effective therapy capable of eliciting durable anti-tumor immunity against NHL and provide a strong rationale for its application in other cancer models that overexpress CD47.

**Fig. 5.**
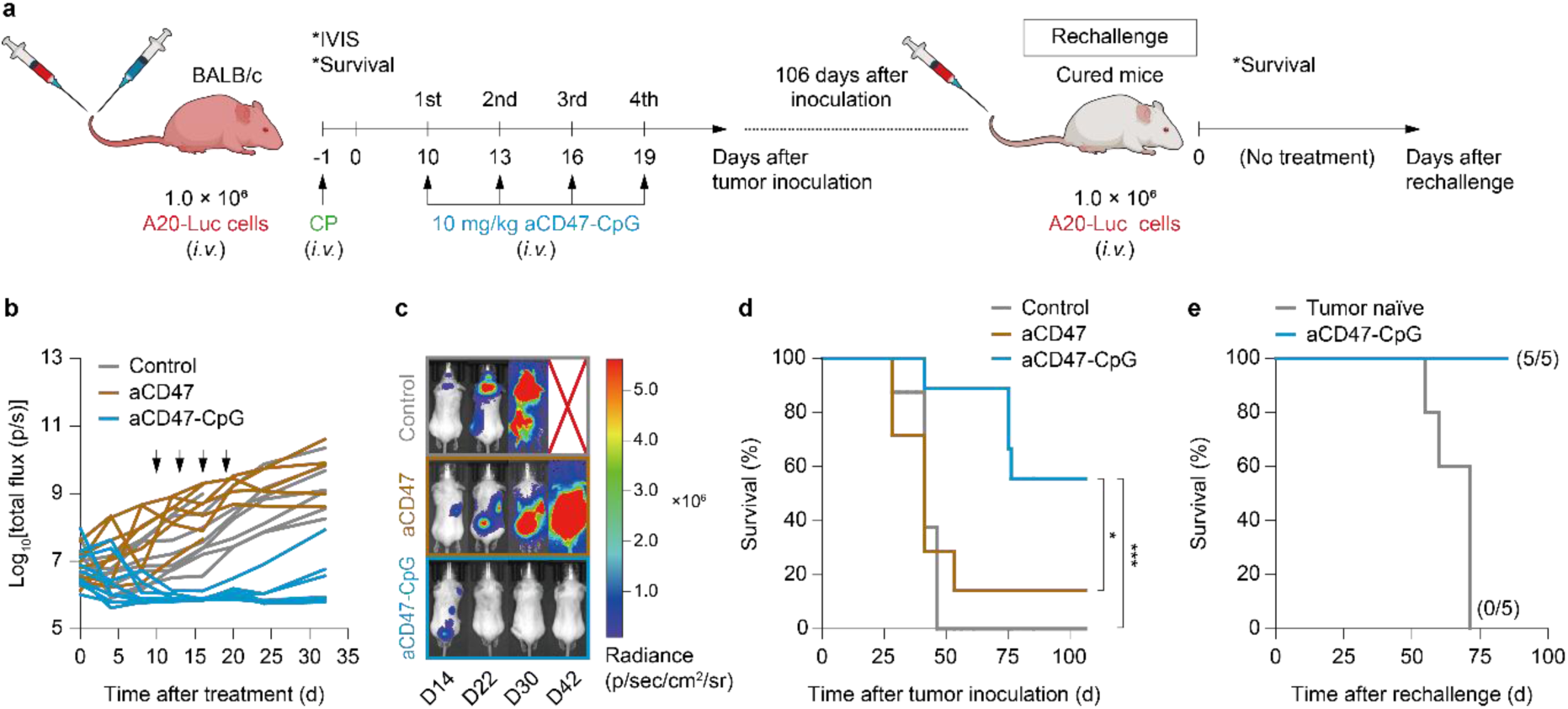
| Systemically administered mouse aCD47-CpG elicits anti-tumor effects in a systemic NHL syngeneic mouse model *in vivo*. **a**, Schematic diagram of tumor inoculation and treatment for a systemic lymphoma model. Following intravenous administration (*i.v.*) of cyclophosphamide (CP), luciferase-labeled A20 (A20-Luc) cells were transplanted *i.v.* into BALB/c mice to establish systemic lymphoma model. Ten days after transplantation, mice were subjected to *i.v.* treatment every third day up to four doses (black arrows) with PBS (control, *n* = 8), aCD47 (*n* = 7, 10 mg/kg), or aCD47-CpG (*n* = 9, 10 mg/kg). For the tumor rechallenge study, the long-term survival animals were rechallenged *i.v.* with the same number of A20-Luc cells 106 days after the initial tumor inoculation. **b**, Quantification of tumor progression for each individual mouse as measured by total flux values acquired via bioluminescence intensity photometry. The black arrows point to administration time. **c**, Representative luciferase imaging of BALB/c mice with systemic tumor receiving PBS, aCD47, or aCD47-CpG in **b**. **d**, Kaplan–Meier survival plot for animals in different treatment groups. **e**, Kaplan– Meier plot depicting the survival of animals following tumor rechallenge. \**p* < 0.05, \*\*\**p* < 0.001 (Log-rank (Mantel-Cox) test).

### Systemically administered aCD47-CpG demonstrates robust anti-tumor activity in syngeneic mouse TNBC models

Building upon our findings in human and mouse NHL, we investigated the efficacy of aCD47 –CpG against TNBC, a highly aggressive and immunosuppressive CD47-overexpressing solid malignancy. Flow cytometry confirmed robust CD47 expression on mouse 4T1 cells (**Extended Data Fig. 5a**), suggesting potential susceptibility to CD47 blockade. In an *in vitro* luminescence-based phagocytosis assay, aCD47-CpG significantly enhanced BMDM-mediated phagocytosis of luciferase-expressing 4T1 cells (**Extended Data Fig. 5b**). Prolonged exposure induced robust upregulation of the co-stimulatory molecule CD40 and elevated secretion of pro-inflammatory cytokines, including TNFα, IL-6, and IL-12, underscoring the immunostimulatory potency against TNBC (**Extended Data Fig. 5c,d**). These effects extended to human TNBC cells where treatment of MDA-MB-231 cells with B6H12-based aCD47-CpG recapitulated key responses, including enhanced phagocytosis, CD40 induction, and pro-inflammatory cytokine release (**Extended Data Fig. 5e–h**). We next examined *in vivo* therapeutic potential of aCD47-CpG and its effect on anti-tumor immune response. To do so, we established an orthotopic syngeneic mouse TNBC model by injecting 4T1 cells, a highly aggressive model characterized by profound immunosuppression and resistance to conventional immunotherapies^35^, into the fourth mammary fat pad of BALB/c mice. Once tumors became palpable, mice received four intravenous injections of PBS, aCD47, or aCD47-CpG (10 mg/kg) every two days (**Fig. 6a**). Treatment with aCD47-CpG markedly suppressed tumor growth compared to control cohorts (**Fig. 6b,c** and **Supplementary Fig. 4**). Remarkably, despite the intrinsically “cold” nature of 4T1 tumors, immune profiling of the TME revealed a substantial enrichment of macrophages, monocytes, and CD8^+^ T cells, alongside with significant reduction of Tregs (**Fig. 6d**). Notably, aCD47-CpG treatment significantly increased the infiltration of both effector and central memory T cells within the tumor (**Fig. 6d**). This enrichment of memory T cell subsets not only suggests the potential for long-term immunological surveillance but also provides cellular-level mechanistic support for the durable immune memory observed in the NHL rechallenge model (**Fig. 5e**). To assess efficacy against metastatic disease, we utilized a TNBC lung metastasis model via intravenous injection of luciferase-expressing 4T1 cells (**Fig. 6e**). BLI analysis revealed a near-complete abrogation of lung metastases in the aCD47-CpG group, characterized by minimal bioluminescent signals relative to controls (**Fig. 6f,g**). Macroscopic analysis of harvested lungs directly confirmed these findings; compared to control treatments, aCD47-CpG markedly reduced the number and size of visible metastatic nodules, indicating a potent inhibitory effect on pulmonary metastasis (**Fig. 6h**). Together, these data demonstrate that aCD47-CpG exerts robust anti-tumor and anti-metastatic effects against TNBC via orchestrating innate and adaptive anti-cancer immunity and reversing the immunosuppressive TME (**Fig. 7**).

**Fig. 6.**
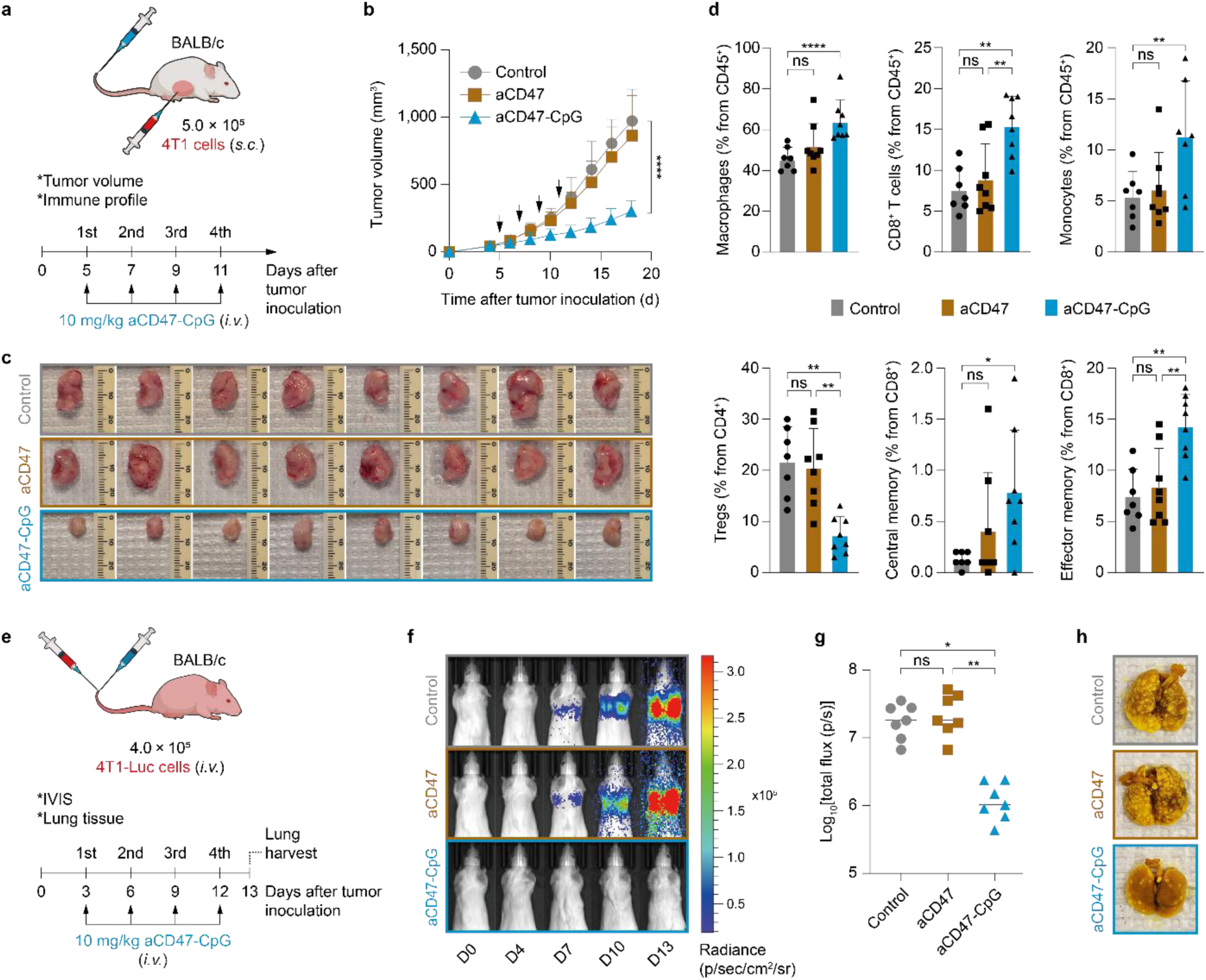
| Systemically administered mouse aCD47-CpG elicits anti-tumor effects in TNBC syngeneic mouse models *in vivo*. **a**, Schematic diagram of tumor inoculation and treatment for an orthotopic TNBC mouse model. Wild-type BALB/c mice were subcutaneously (*s.c.*) implanted with 4T1 cells into the fourth right mammary fat pad. Five days after tumor inoculation, aCD47-CpG was given via intravenous administration (*i.v.*) at 10 mg/kg every two days up to four doses (black arrows). For comparison, animals in different groups received either PBS (control) or aCD47 following the identical dosing schedule (n = 8 per group). **b**, Tumor growth of mice treated in **a**. Tumor progression was monitored by tumor volume measurement using a caliper. **c**, Image of excised 4T1 TNBC tumors from mice. **d**, Flow cytometry analysis of immune cell profile in tumor microenvironment, including myeloid and lymphoid population. **e**, Schematic diagram of tumor inoculation and treatment for a metastatic TNBC mouse model. Luciferase-expressing 4T1 (4T1-Luc) cells were transplanted *i.v.* into BALB/c mice to establish a metastatic TNBC model. Three days after transplantation, mice were subjected to *i.v.* administration every third day up to four doses with PBS (control), aCD47 (10 mg/kg), or aCD47-CpG (10 mg/kg) (*n* = 7 per group). One day after the final dose, the lungs were harvested, and metastases were imaged. **f**, Representative luciferase imaging of BALB/c mice with metastatic tumors receiving PBS, aCD47, or aCD47-CpG. **g**, Quantification of lung tumor metastasis for each individual mouse as measured by flux values acquired from **f** via bioluminescence intensity photometry. **h**, Representative images of harvested lungs from mice in each treatment group. **b**, \*\*\*\**p* < 0.0001 (two-way ANOVA with Tukey’s multiple comparisons test). **d**,**g**, ns: non-significant, \**p* < 0.05, \*\**p* < 0.01, \*\*\*\**p* < 0.0001 (one-way ANOVA with Tukey’s multiple comparisons test).

**Fig. 7.**
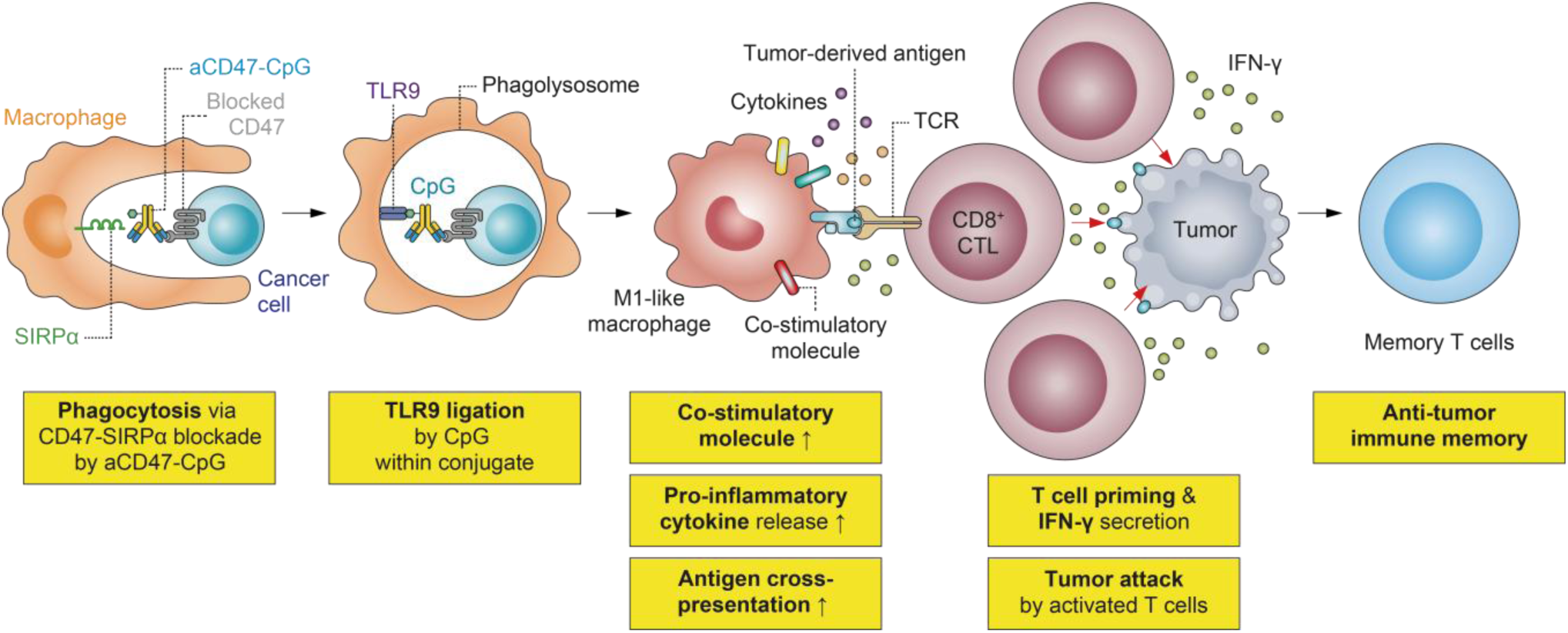
| Experimentally validated anti-tumor mechanism-of-action of aCD47-CpG. The aCD47-CpG functions through a multi-step process to enhance anti-tumor immunity. Initially, the anti-CD47 antibody component of the conjugate blocks the interaction of CD47, “don’t eat me” signal, expressed on cancer cells with SIRPα on macrophages, allowing macrophages to recognize and phagocytose tumor cells. Upon engulfing the cancer cells, CpG piggybacked on the conjugate activates TLR9 signaling within the phagolysosome, leading to potent immune activation responses, such as upregulation of co-stimulatory molecules, release of pro-inflammatory cytokines, and enhancement of antigen cross-presentation. These steps collectively enable effective priming of CD8^+^ cytotoxic T lymphocytes (CTLs), which secrete interferon gamma (IFN-γ), further amplifying the anti-tumor immune response and potentially recruiting additional immune cells to the tumor site. After activated CD8^+^ CTLs destroy the tumor cells, memory T cells are generated, conferring long-lasting immunity against tumor recurrence.

## Discussion

Here, we demonstrate that a TLR9 agonist CpG conjugated to an aCD47 bridges initial innate phagocytosis to sustained adaptive anti-tumor immunity. Our data show that aCD47-CpG functions through a multi-step and spatiotemporally coordinated process commenced with aCD47-mediated tumor cell engulfment, followed by phagolysosomal CpG delivery and TLR9 stimulation. This initial cascade triggers macrophage activation and repolarization to deter tumor growth as revealed in our *in vivo* study with a human NHL xenograft model. Furthermore, the reprogrammed macrophages drive robust antigen cross-presentation, thereby effectively priming tumor-specific T cells. This immunological remodeling translates into potent primary tumor control and resistance to recurrence and metastasis, respectively, in fully immunocompetent syngeneic NHL and TNBC models *in vivo*, ultimately culminating in durable anti-tumor immune memory.

The ability to elicit such memory represents the most critical therapeutic advantage of the aCD47-CpG platform. Remarkably, mice with systemic NHL that experienced complete tumor remission following aCD47-CpG treatment resisted subsequent tumor rechallenge. This functional protection was in good agreement with cellular profiling of the TNBC TME, revealing an expansion of effector and central memory T cell populations upon aCD47-CpG treatment. Mechanistically, the establishment of this sustained immunity is associated with the remodeling of the immunosuppressive TME. Our data demonstrate that aCD47-CpG promotes M2-to-M1 macrophage repolarization and intratumoral cytotoxic CD8^+^ T-cell infiltration while reducing Tregs in TME. Of note, Tregs suppress tumor-specific CD8^+^ T cell activation and expansion^36^, and the magnitude of this initial effector response dictates the size of the subsequently generated memory T cell pool^37^. To this end, mitigation of Treg-mediated immunosuppression is a prerequisite for durable T cell immunity. Furthermore, the strong IFN-γ secretion driven by aCD47-CpG *in vitro* indicates its ability to promote a Th1-polarized environment, which is essential for acute tumoricidal activity and the expansion of effector T cells^38^. Therefore, the aCD47-CpG-mediated shift toward an immune-permissive TME likely has contributed to both immediate tumor clearance and long-term immune surveillance.

At the cellular level, this immune remodeling is driven by sustained macrophage activation that extends beyond mere facilitation of tumor cell phagocytosis by CD47 blockade. Our data showed that aCD47-CpG exhibited an initial delay in phagocytosis, likely reflecting its attenuated CD47-binding affinity, but the phagocytic activity surpassed unconjugated aCD47 at later time points. This functional reversal indicates that TLR9 signaling, triggered by the internalization of aCD47-CpG and subsequent target engagement, drives durable phagocytosis and anti-tumor immunomodulation. The concept of utilizing aCD47-mediated internalization of immune-stimulating agents has been pioneered by a recently introduced aCD47-listeriolysin O (LLO) conjugate^39^. This system works by LLO forming pores on phagolysosomal membranes to release the engulfed tumor components into the cytosol for immune sensing. Notably, our aCD47-CpG platform offers a distinct spatiotemporal feature by enabling the target engagement of CpG in endosomal and phagolysosomal compartments upon internalization without necessity of cytosolic release. This spatially coordinated signaling drives effective macrophage polarization toward an anti-tumor M1-like state, as evidenced by the upregulation of iNOS and co-stimulatory molecules, including CD80, CD86, and CD40. This targeted delivery of immune-stimulating payloads presumably facilitated the innate-to-adaptive immune transition to create robust and durable *in vivo* anti-tumor efficacy observed against multiple cancer models.

Recent advancements in ISACs have leveraged tumor-specific antibodies, such as those targeting HER2, EGFR, or CD22, to deliver innate immune agonists (*e.g.*, TLR7/8, TLR9, and STING agonists) to the TME^20, 40–43^. Mechanistically, these conventional antigen-specific ISACs depend on a tripartite mechanism, comprising the simultaneous coordination of tumor antigen recognition, payload – mediated innate immune sensing, and FcγR engagement, to drive myeloid cell internalization and phagocytosis^20, 42, 43^. One limitation is that their clinical translation is inherently constrained by tumor antigen heterogeneity and the stringent requirement for robust target expression within specific cancer indications^41, 43^. On the other hand, aCD47-CpG may address this antigen restriction by exploiting a ubiquitous, pan-cancer myeloid phagocytosis checkpoint. The blockade of the CD47-SIRPα axis intrinsically lowers the phagocytic threshold, and thus aCD47-CpG does not require the activation of FcγR engagement to initiate engulfment. This is evidenced by the potent anti-tumor immunity achieved against multiple preclinical cancer models by the treatment with aCD47-CpG based on a mouse IgG1 backbone that possesses minimal affinity for activating FcγRs^44^. Therefore, by superseding the FcγR dependency inherent to tumor-specific ISACs, the aCD47-CpG platform offers distinct mechanistic versatility. In addition, by enabling the accommodation of an Fc-silent backbone, aCD47-CpG potentially mitigates systemic Fc-mediated toxicities for its application to a broad cancer patient population.

Securing clinical safety is essential to the development of CD47-targeted therapies. Initial clinical trials evaluating an investigational CD47, magrolimab, across various advanced solid tumors and hematologic malignancies, including breast cancer and lymphoma, demonstrated manageable safety profiles without dose-limiting toxicities^45, 46^. In parallel, systemic intravenous administration of CpG has been shown to be well-tolerated by patients with previously treated NHL, exhibiting minimal adverse events^47^. Consistent with the observed clinical tolerability of these individual components, our aCD47-CpG conjugate exhibited acceptable *in vivo* safety in our preclinical studies. However, broader clinical experience has revealed that the safety profile of CD47 blockade varies considerably depending on cancer type and combination regimen^26, 48^. For instance, a recent phase 3 study in acute myeloid leukemia evaluating magrolimab in combination with venetoclax and azacitidine reported an unfavorable risk-to-benefit profile^48^. In contrast, an earlier clinical trial in NHL demonstrated that magrolimab combined with rituximab yielded promising therapeutic activity without clinically significant safety events^26^. These clinical observations collectively underscore that the safety of CD47-targeted agents must be carefully evaluated within specific therapeutic contexts for their clinical implementation.

Overall, we demonstrate here that the marriage of innate tumor cell engulfment with adaptive immune priming via aCD47-CpG enhances the therapeutic potential and coverage of CD47-targeted therapies. This ISAC platform uniquely enables the decoupling of therapeutic efficacy from specific tumor antigen and FcγR dependencies while maintaining systemic safety to potentially benefit a broad spectrum of cancer patient populations upon clinical development.

## Methods

### Cell lines and cell culture

Mouse A20 lymphoma cells (TIB-208) and E.G7-OVA cells (CRL-2113) were purchased from ATCC. A20 cells expressing GFP and luciferase (A20-GFP/Luc) were a kind gift from Dr. Robert Zeiser (University of Freiburg). Mouse 4T1 breast cancer cell line expressing yellow fluorescent protein (YFP) and luciferase (4T1-YFP/Luc), human Raji lymphoma cells expressing GFP and luciferase (Raji-GFP/Luc), and human MDA-MB-231 cells expressing GFP and luciferase (MDA-MB-231-GFP/Luc) were obtained from Translational Laboratory Shared Resources (TLSR) of University of Maryland School of Medicine. These cell lines were validated by short tandem repeat analysis and the mouse antibody production test. Tumor cell lines were cultured at 37°C and 5% CO_2_ in Roswell Park Memorial Institute (RPMI) 1640 medium (Gibco, Thermo Fisher Scientific, 11875-093) supplemented with 10% (v/v) fetal bovine serum (FBS; GeminiBio, 100-106H-500), 100 U/mL of penicillin, and 100 µg/mL of streptomycin (Pen-Strep; Gibco, Thermo Fisher Scientific, 15140122) (complete RPMI). The culture media for the A20, A20-GFP/Luc, and E.G7-OVA cell lines were additionally supplemented with 0.05 mM 2-mercaptoethanol (Sigma-Aldrich, M3148). Cells were split twice per week and counted using a hemacytometer after staining with trypan blue.

### Mice

Six-to eight-week-old female BALB/c, BALB/c nude, and C57BL/6 mice were purchased from Charles River Laboratories. Transgenic OT-I mice (C57BL/6-Tg(TcraTcrb)1100Mjb/J) were obtained from The Jackson Laboratory and bred in-house. All animals were housed at the animal facility of University of Maryland School of Medicine. All mouse experiments were conducted according to the protocols approved by the Institutional Animal Care and Use Committee (IACUC) of the University of Maryland School of Medicine.

### aCD47-CpG production and characterization

For chemical conjugation, anti-mouse CD47 (aCD47; clone MIAP410, Bio X Cell, BE0283) or anti-human CD47 antibody (clone B6H12, Bio X Cell, BE0019-1), buffer-exchanged into phosphate-buffered saline (PBS) supplemented with 10 mM EDTA (PBSE; pH 7.4) using a PBSE-equilibrated PD-10 desalting column (Cytiva, 17085101), was mixed with the heterobifunctional crosslinker Sulfo-SMCC (Thermo Fisher Scientific, 22322) dissolved in distilled water. The mixture was incubated with continuous rotation at room temperature for 1 h according to the manufacturer’s protocol and then the excess linker was removed using a PD-10 column. Thiol-modified CpG (10 mg/mL, Integrated DNA Technologies) was reduced with an excess of dithiothreitol (DTT; Sigma-Aldrich, D0632) at 100 mM in PBSE for 2 h at room temperature and protected from light. After removal of excess DTT using a PBSE-equilibrated PD-10 column, the maleimide-activated antibodies were combined with the reduced CpG at molar ratio of 1:20, which was incubated at 4°C overnight. Unconjugated CpG was removed from the mixture by size exclusion chromatography (SEC) on a Superose™ 6 Increase 10/300 GL column (Cytiva, 29091596) equilibrated with PBS. The purity and conjugation pattern of the resulting aCD47-CpG were confirmed by Coomassie blue staining for aCD47 and SYBR Gold staining for CpG in SDS-PAGE analysis. The concentration and CpG-to-aCD47 ratio (*i.e.*, drug-antibody ratio) were determined by measuring the absorbance of both molecules at 260 and 280 nm on a NanoDrop One (Thermo Fisher Scientific), as described elsewhere^49^. The purified conjugates were filtered through a 0.22-µm sterile filter and stored at 4°C until use.

### ELISA

The binding specificities of anti-CD47 antibody and CpG within aCD47-CpG to CD47 and TLR9 were determined by ELISA using recombinant mouse CD47 (mCD47; SinoBiological, 57231-M08H) and mouse TLR9 protein (mTLR9; LifeSpan BioSciences, LS-G24092), respectively. All steps were conducted at room temperature unless otherwise noted. Nickel-coated 96-well plates (Pierce, Thermo Fisher Scientific, 15442) were coated with 100 µL of 1 µg/mL mCD47 or mTLR9 dissolved in tris-buffered saline (TBS, pH 7.4) for 2 h. The plates were washed three times with 200 µL of TBS supplemented with 0.1% (v/v) Tween 20 (Sigma-Aldrich, P7949) (TBST) and blocked for 1 h with 100 µL of PBS containing 1% (w/v) bovine serum albumin (BSA; Sigma-Aldrich, A7906) filtered through a 0.22-µm membrane. After washing with TBST three times, serially diluted aCD47-CpGs or isotype control IgG (100 µL) were added to the plates and incubated at 4°C overnight. Following TBST washing step, the plates were treated for 1 h with 100 µL of goat anti-mouse IgG antibody conjugated with horseradish peroxidase (HRP) (Invitrogen, Thermo Fisher Scientific, 31430; 1:5,000). The plates were washed three times with TBST and 100 µL of 1-Step^TM^ Slow TMB (Thermo Fisher Scientific, 34024) was added to each well. After color was developed for 10 min, 100 μL of 1 N sulfuric acid (Fisher Scientific, Thermo Fisher Scientific, SA212-1) was added to the plates and then the absorbance at 450 nm was measured using a Synergy H1 Plate Reader (BioTek). K_D_ values were obtained using Graph Pad Prism 8 software.

### NF-κB reporter assay

RAW-Blue^TM^ cells, engineered to monitor the NF-κB and AP-1 response upon TLR9 stimulation, were purchased from InvivoGen (raw-sp) and cultured in Dulbecco’s Modified Eagle Medium (DMEM) supplemented with 10% (v/v) FBS, 1× Pen-Strep, and 100 µg/mL of Normocin™ according to the manufacturer’s protocol. Briefly, 1 × 10^5^ cells seeded in 96-well plate were treated with CpG at a concentration of 1 µg/mL in the presence or absence of A20 cells and incubated at 37°C for 18–24 h. Secreted alkaline phosphatase (SEAP) activity was confirmed by measuring the absorbance at 620 nm using a microplate reader (Synergy H1 Plate Reader) after mixing 20 µL of cell culture supernatant with 180 µL of QUANTI-Blue solution followed by incubation at 37°C for 2 h.

### Generation of bone marrow-derived macrophages (BMDMs)

Hematopoietic stem cells from the tibias and femurs of 6– to 8-week-old BALB/c and C57BL/6 mice were flushed using a syringe into complete RPMI. After gently pipetting, cells were filtered through a 40-µm cell strainer (Thermo Fisher Scientific, 352340) and centrifuged at 4°C for 10 min at 450 × *g*. The RBC population was lysed by resuspending the cell pellet in 1× RBC lysis buffer (BioLegend, 420301), followed by incubation at 4°C for 5 min. After quenching with a large volume of cold PBS, cells were centrifuged and resuspended in complete RPMI supplemented with 20 ng/mL macrophage-colony stimulating factor (M-CSF; PeproTech, 315-02). The cell suspension was plated on 100-mm petri dishes and cultured for 7 days with M-CSF added every other day. Differentiation into BMDMs was confirmed by flow cytometry using APC-labeled anti-mouse F4/80 antibody (Invitrogen, Thermo Fisher Scientific, 17-4801-82).

### *In vitro* phagocytosis assay

For *in vitro* phagocytosis, tumor cells virally expressing GFP or YFP (3 × 10^4^ cells) were pre-incubated with indicated concentrations of control, aCD47, or aCD47-CpG for 30–60 min at 4°C in 96-well round-bottom ultra-low attachment microplate (Corning, 7007). BMDMs were washed with PBS three times and incubated with trypsin-EDTA (Gibco, Thermo Fisher Scientific, 25200072) for up to 15 min at 37°C followed by mild scraping. After adding complete RPMI to inactivate the trypsin, BMDMs were centrifuged at 400 × *g* for 5 min and resuspended in serum-free RPMI containing anti-mouse CD16/32 blocking antibodies (1:50 dilution) (BioLegend, 101301) followed by incubation at 4°C for 20 min. BMDMs were washed with PBS and then added to the plate (6 × 10^4^ cells), which was incubated at 37°C for 2 h. Phagocytosis images were obtained by phase contrast microscopy (ECHO Revolve microscope, RVL2-K) using the ECHO Pro software. For the flow-based phagocytosis assay, after staining BMDMs with APC-labeled anti-F4/80 antibodies (1:200 dilution in serum-free RPMI), data were collected by flow cytometry (Cytek® Aurora Cytometer) using SpectroFlo version 3.3.0 and analyzed using FlowJo software version 10.10.0. Phagocytosis was calculated by the percentage of BMDMs which phagocytosed tumor cells (*i.e.*, the ratio of GFP/YFP^+^APC^+^ cells to APC^+^ cells).

### Luminescence-based long-term macrophage killing (LB-LTMK) assay

The LB-LTMK assay followed a previously described method^27^. Briefly, cancer cells expressing luciferase (3 × 10^4^ cells) were co-cultured with BMDMs (6 × 10^4^ cells) prepared as in the *in vitro* phagocytosis assay at 37°C for 24 h in 96-well cell culture plates (Corning, 353075) using complete RPMI in the presence of control, aCD47, or aCD47-CpG with a total volume of 150 µL per well. For *in vitro* luciferase assay, cells were lysed by adding Reporter Lysis 5× Buffer (Promega, E4030). Plates were placed at −80°C for 30 min and then moved to 37°C for 30 min for complete cell lysis. The luminescence signal was detected in Turner BioSystems 20/20n Luminometer after mixing 10 µL of cell lysate with 100 µL of Luciferase Assay Substrate (Promega, E1501). The phagocytosis was calculated as the ratio of relative luminescence signals in the treatment group to the signals in the cancer cell-only group. For the kinetics of phagocytosis, the same procedure was performed at each indicated time point.

### Cellular activation assays

Cellular activation was investigated through cell surface expression of activation markers and cytokine levels secreted into the supernatant. Tumor cells were plated and treated with control, aCD47, or aCD47-CpG and then BMDMs were added as described in LB-LTMK assay. After a 24-h incubation, cells were stained for surface markers using antibodies against mouse CD80 (BioLegend, 104713), CD86 (BioLegend, 105021), and CD40 (BioLegend, 124609). For intracellular iNOS detection, cells were fixed and permeabilized using a Foxp3/Transcription Factor Staining Buffer Set (Invitrogen, 00-5523-00) according to manufacturer’s instructions prior to staining with an anti-iNOS antibody (BioLegend, 696807). All antibodies were diluted 1:200 in FACS buffer comprising PBS supplemented with 2% (v/v) FBS and 2 mM EDTA. Data were acquired on an LSR II flow cytometer (BD Biosciences) utilizing FACSDiva software (v8.0.1) and analyzed using FlowJo software (v10.10.0). For cytokine level analysis, TNFα (R&D Systems, DY410), IL-6 (R&D Systems, DY406), and IL-12 (R&D Systems, DY499) levels in cell culture supernatants were tested using their respective ELISA kits according to the manufacturer’s instructions.

### Antigen cross-presentation

E.G7-OVA cells were plated and treated with control, aCD47, or aCD47-CpG and then BMDMs were added as described in the LB-LTMK assay. After 24 h of incubation, cells were stained at 4°C for 1 h with FITC-labeled anti-mouse F4/80 (diluted 1:200, BioLegend, 123107) and APC-labeled anti-SIINFEKL-H2Kb antibodies (diluted 1:200, Invitrogen, 17-5743-82) in the dark. Cross-presentation of the ovalbumin-derived SIINFEKL peptide bound to H-2Kb of MHC class I was examined by flow cytometry (BD Biosciences, LSR II) using FACSDiva version 8.0.1 and analyzed using FlowJo software version 10.10.0. The level of cross-presentation was quantified as the mean fluorescence intensity of APC within the FITC^+^ cell population.

### CD8^+^ T cell isolation

Transgenic OT-I mice underwent cardiac perfusion with PBS, and then spleens were carefully collected, washed with Hank’s Balanced Salt Solution (HBSS) containing 2% (v/v) FBS, and dissociated through 40-µm cell strainer (Thermo Fisher Scientific, 352340) using mechanical force. From this step onward, all procedures were conducted at 4°C under sterile conditions. After centrifugation at 400 × *g* for 5 min, the RBCs were lysed by resuspending the cell pellet in 1× RBC lysis buffer (BioLegend, 420301), followed by incubation for 5 min. After quenching with large volume of cold PBS, cells were centrifuged, resuspended in HBSS supplemented with 2% FBS, and filtered with 40-μm cell strainer for removal of cell aggregation. CD8^+^ T cells were isolated from the total cell population using MojoSort^TM^ mouse CD8 T Cell Isolation Kit (BioLegend, 480008) according to the manufacturer’s instructions. Briefly, 10 µL of the Biotin-Antibody cocktail for depleting non CD8^+^ T cells was added to 100 µL of 1 × 10^8^ cells/mL cell suspension, which was incubated on ice for 15 min. Streptavidin Nanobeads were added to the tube and then incubated on ice for 15 min. After adding 1 mL of PBS and subsequently placing the tube in the magnet stand for 5 min, CD8^+^ T cells in the liquid were collected. This procedure was performed twice. Following centrifugation at 400 × *g* for 5 min, cells were suspended in FACS buffer, and labeled with 1.5 µM of CellTrace^TM^ CFSE Cell Proliferation kit (Thermo Fisher Scientific, C34554) according to the manufacturer’s instructions.

### T cell priming

For *in vitro* T cell priming, E.G7-OVA cells were plated and treated with control, aCD47, or aCD47-CpG and then BMDMs were added as described in LB-LTMK assay. After 24 h of incubation, CD8^+^ T cells labeled with CFSE and isolated from OT-I mouse spleens were added to the plates at a T cell-to-BMDMs ratio of 10:1. After 3 days, cells were incubated with complete RPMI containing anti-mouse CD16/32 blocking antibodies (1:50 dilution) at 4°C for 20 min followed by staining with Brilliant Violet 510^TM^ (BV510) anti-mouse CD8a antibodies (1:100 dilution) (BioLegend, 100751) for flow cytometry analysis at 4°C for 1 h. T cell proliferation was assessed by monitoring CFSE dilution within BV510 ^+^ cells using a FACS Canto II (BD Biosciences) equipped with FACSDiva software, with subsequent data analysis using FlowJo software (version 10.10.0). IFN-γ secreted into the cell culture supernatant following T cell priming was detected using Mouse IFN-γ ELISA Kit (Invitrogen, KMC4021) according to the manufacturer’s instructions.

### *In vivo* toxicology study

For toxicology study, the clinically validated low-dose priming strategy was employed where healthy BALB/c mice were initially treated intravenously (*i.v.*) with a priming dose of 1 mg/kg, followed two days later by three escalating doses of aCD47-CpG at 5, 10, or 20 mg/kg. MTD was established by monitoring survival and weight loss of mice. For the hematotoxicity study, healthy BALB/c mice were injected *i.v.* with a single dose of aCD47-CpG at 5, 10, or 20 mg/kg without priming dose and blood was collected at the indicated time points for complete blood count analysis.

### *In vivo* biodistribution and tumor specificity analysis

To evaluate the tumor-targeting capability and biodistribution profile of the aCD47-CpG conjugate, a Cy5.5-labeled CpG was conjugated to the anti-CD47 antibody. Subsequently, a syngeneic NHL model was established by subcutaneously implanting A20 lymphoma cells (2.5 × 10^6^) into the right flank of immunocompetent BALB/c mice. When the average tumor volume reached approximately 100–200 mm³, tumor-bearing mice were randomized and intravenously administered the Cy5.5-labeled aCD47-CpG conjugate via tail vein at a dose of 10 mg/kg (MTD). To verify CD47-mediated tumor specificity, a Cy5.5-labeled isotype IgG-CpG conjugate at an equivalent dose, as well as PBS, were included as controls. At 24-h post-injection, mice were euthanized, and tumors along with major organs (brain, heart, lung, liver, spleen, and kidney) were excised. The fluorescence intensity of the harvested tissues was quantified *ex vivo* to assess biodistribution, and tumor specificity was further analyzed by comparing the fluorescence intensities of the tumors across the groups.

### *In vivo* anti-cancer efficacy study in a human lymphoma xenograft model

To evaluate the therapeutic efficacy of aCD47-CpG against human tumors, an *in vivo* xenograft model was established. BALB/c nude mice were inoculated with a subcutaneous injection of 5.0 × 10^6^ human Raji lymphoma cells into the right flank. When tumors reached a palpable size, animals were randomly assigned to different groups and treated intravenously with PBS, aCD47, or aCD47-CpG at a dose of 10 mg/kg every three days for a total of four doses. To validate the macrophage-dependent mechanism of action, a separate cohort received an intraperitoneal injection of macrophage-depleting clodronate liposomes (LIPOSOMA) three times a week, starting one day prior to the initial aCD47-CpG treatment. Tumor progression was monitored by caliper measurement using the formula (length × width^2^)/2. At the study endpoint, tumors were excised, weighed, and processed for flow cytometry analysis of the tumor microenvironment.

### *In vivo* anti-cancer efficacy study in a systemic mouse lymphoma syngeneic model

For the *in vivo* anti-cancer efficacy study in NHL, a systemic disseminated lymphoma mouse model was established. BALB/c mice were inoculated with intravenous injection of 1 × 10^6^ A20-Luc cells. Following the confirmation of systemic lymphoma development by live-animal bioluminescence imaging, animals were randomly assigned to three different groups and systemically treated with PBS, aCD47, or aCD47-CpG at the MTD every third day up to four doses. For the tumor rechallenge study, long-term survivors were rechallenged *i.v.* with A20-Luc cells of same cell number 106 days after the initial tumor inoculation. As a control, age-matched naïve BALB/c mice were challenged with the same tumor cells. Animals were then monitored for survival and Kaplan–Meier survival comparison was performed using log-rank (Mantel-Cox) test in Graph Pad Prism 8 software.

### *In vivo* anti-cancer efficacy study in mouse breast cancer syngeneic model

For *in vivo* anti-cancer efficacy study in TNBC, two complementary mouse models of TNBC were established. For an orthotopic TNBC mouse model, 5 × 10^5^ 4T1 cells were injected *s.c.* into the fourth right mammary fat pad of BALB/c mice. Tumors were grown to reach approximately 50 mm^3^ in volume and then aCD47-CpG was given via systemic administration at the MTD every third day up to four doses. For comparison, animals in different groups received either PBS or aCD47 following the identical dosing schedule. Throughout the course of the treatments and afterwards, the tumor volume was measured with caliper using the formula (length × width^2^)/2 and tumor tissues were harvested for immune cell profiling with flow cytometry analysis. For a metastatic TNBC mouse model, 4T1-Luc cells (4 × 10^5^) were injected *i.v.* into the tail vein of BALB/c mice. After 3 days, the mice were randomly assigned into three groups and systemically treated with PBS, aCD47, or aCD47-CpG at the MTD every third day up to four doses. Tumor metastasis progression was measured by live-animal bioluminescence imaging. At the end of the study, the lungs were harvested and stained with Bouin’s solution (Ricca Chemical, 112016) at room temperature for 24 h. The following day, the organs were immersed in 70% ethanol for storage, and metastatic nodules were macroscopically evaluated.

### Flow-based immune cell profiling of tumor tissues

After euthanizing mice, tumors were excised and minced into small pieces (1–3 mm) in a petri dish with a scalpel. The minced tumor tissues were incubated with 1 mg/mL collagenase solution formulated with collagenase D (Roche, 11088882001) in serum-free RPMI for 1 h at 37°C with gentle shaking every 10 min. The digested tissues were mechanically dissociated through a 40-µm cell strainer using a syringe plunger and the dissociated single cell suspension was centrifuged at 400 × *g* for 5 min at 4°C. The RBCs in the cell pellet were lysed by resuspension in 1× RBC lysis buffer (BioLegend, 420301), followed by incubation at 4°C for 5 min. After quenching with large volume of cold PBS, cells were centrifuged and resuspended in 40% (v/v) Percoll solution (Cytiva, 17089101). After layering 80% (v/v) Percoll solution under 40% Percoll with a glass Pasteur pipet, the tubes were centrifuged at 600 × *g* for 20 min at 4°C with low acceleration and no brake. Immune cell layers were carefully transferred into a new 15-mL tube and then centrifuged again for removal of residual Percoll. The cell pellets were resuspended in 50 µL of FACS buffer containing anti-mouse CD16/32 blocking antibodies (1:50 dilution) at 4°C for 20 min. Subsequently, cells were stained at 4°C for 30 min in the dark using antibody cocktails designed to profile specific immune cell subsets. For generalized immune cell profiling, the cocktail consisting of the following fluorescence-conjugated antibodies specific for representative markers of each immune cell subset: AF700 anti-mouse CD45.2 antibody (BioLegend, 109822), PE-Dazzle594 anti-mouse CD11b antibody (1:400 dilution) (BioLegend, 101256), BV711 anti-mouse CD3 antibody (BioLegend, 100241), FITC anti-mouse F4/80 antibody (Invitrogen, 11-4801-82), BV421 anti-mouse Ly6G antibody (BioLegend, 127628), PE-Cy7 anti-mouse Ly6C antibody (1:400 dilution) (BioLegend, 128018), eFluor 506 anti-mouse CD4 antibody (Invitrogen, 69-0042-82), PE-Cy7 anti-mouse CD8a antibody (BioLegend, 100722), BV605 anti-mouse CD25 antibody (BioLegend, 102036), PerCP-Cy5.5 anti-mouse CD44 antibody (BioLegend, 103032), and BV421 anti-mouse CD62L antibody (1:200 dilution) (BioLegend, 104436). For detailed myeloid cell profiling, particularly to evaluate macrophage polarization in xenograft tumor models, an additional flow cytometry panel was utilized. This cocktail included AF700 anti-mouse CD45.2 (BioLegend, 109822), BUV395 anti-mouse CD11b (Invitrogen, 363-0112-82), BV711 anti-mouse CD3 (BioLegend, 100241), FITC anti-mouse F4/80 (Invitrogen, 11-4801-82), BV421 anti-mouse Ly6G antibody (BioLegend, 127628), PE-Cy7 anti-mouse Ly6C antibody (BioLegend, 128018), BV510 anti-mouse MHCII (BioLegend, 107635), and PE anti-mouse CD11c (BioLegend, 117307). Unless otherwise specified, all antibodies were used at a dilution of 1:100. For intracellular Foxp3, iNOS, and EGR2 staining, fixation and permeabilization steps were additionally included using the Foxp3/Transcription Factor Staining Buffer Set (Invitrogen, 00-5523-00) according to the manufacturer’s instructions, followed by staining with FITC anti-mouse Foxp3 antibody (Invitrogen, Thermo Fisher Scientific, 11-5773-82), AF594 anti-mouse iNOS antibody (BioLegend, 696803), and PE anti-mouse EGR2 antibody (Invitrogen, 12-6691-82). Immune cell profiling in tumor tissues were assessed by flow cytometry (Cytek® Aurora Cytometer) using SpectroFlo version 3.3.0 and analyzed using FlowJo software version 10.10.0.

## Statistical analysis

Statistical analysis was conducted using one-way (one independent variable) or two-way ANOVA (two or more independent variables) with Tukey’s multiple comparisons test to compare 3 or more groups (GraphPad Prism software version 8.0). All statistical tests were two-sided. All data are plotted as mean ± standard deviation. For the statistical test, asterisks are used to indicate statistical significance (\**p <* 0.05, \*\**p <* 0.01, \*\*\**p <* 0.001, \*\*\*\**p <* 0.0001), non-significant values (*p>*0.05) are denoted as n.s.

## Data availability

The data supporting the findings of this study are available within the article and its Supplementary Information. Additional materials are available from the corresponding author upon reasonable request.

## Acknowledgements

This work was supported by the National Institutes of Health (R01NS119609 to J.S.S.), the Maryland Innovation Initiative (MII) of the Maryland Technology Development Corporation (TEDCO) (0722-005 to J.S.S.), and the Sejong Science Fellowship from the National Research Foundation of Korea (RS-2024-00358493 to B.K.). The authors thank the Flow Cytometry Shared Service of the University of Maryland Marlene and Stewart Greenebaum Comprehensive Cancer Center for access to instrumentation.

## Author information

### Authors and Affiliations

**Department of Neurosurgery, University of Maryland School of Medicine, Baltimore, MD, USA**

B. Kong, D. Lee, G. Kwak & J. S. Suk

**University of Maryland Medicine Institute for Neuroscience Discovery (UM-MIND), University of Maryland School of Medicine, Baltimore, MD, USA**

B. Kong, D. Lee, G. Kwak & J. S. Suk

**Department of Ophthalmology, Johns Hopkins University School of Medicine, Baltimore, MD, USA**

S. W. Chung

**Department of Neurosurgery, Johns Hopkins University School of Medicine, Baltimore, MD, USA**

J. S. Suk

**Fischell Department of Bioengineering, University of Maryland, College Park, MD, USA**

J. S. Suk

### Contributions

B.K. designed and performed the experiments, analyzed the data, and wrote the manuscript. S.W.C. provided conceptual input and discussed the results. D.L. and G.K. assisted with the *in vivo* animal experiments and discussed the results. J.S.S. supervised the project, acquired funding, and discussed the results. All authors reviewed and approved the final manuscript.

### Corresponding author

Correspondence to Jung Soo Suk.

## Ethics declarations

### Competing Interests

J.S.S. and S.W.C. are inventors on a patent application related to the aCD47-CpG conjugate technology described in this manuscript. The remaining authors declare no competing interests.

**Extended Data Fig. 1.**
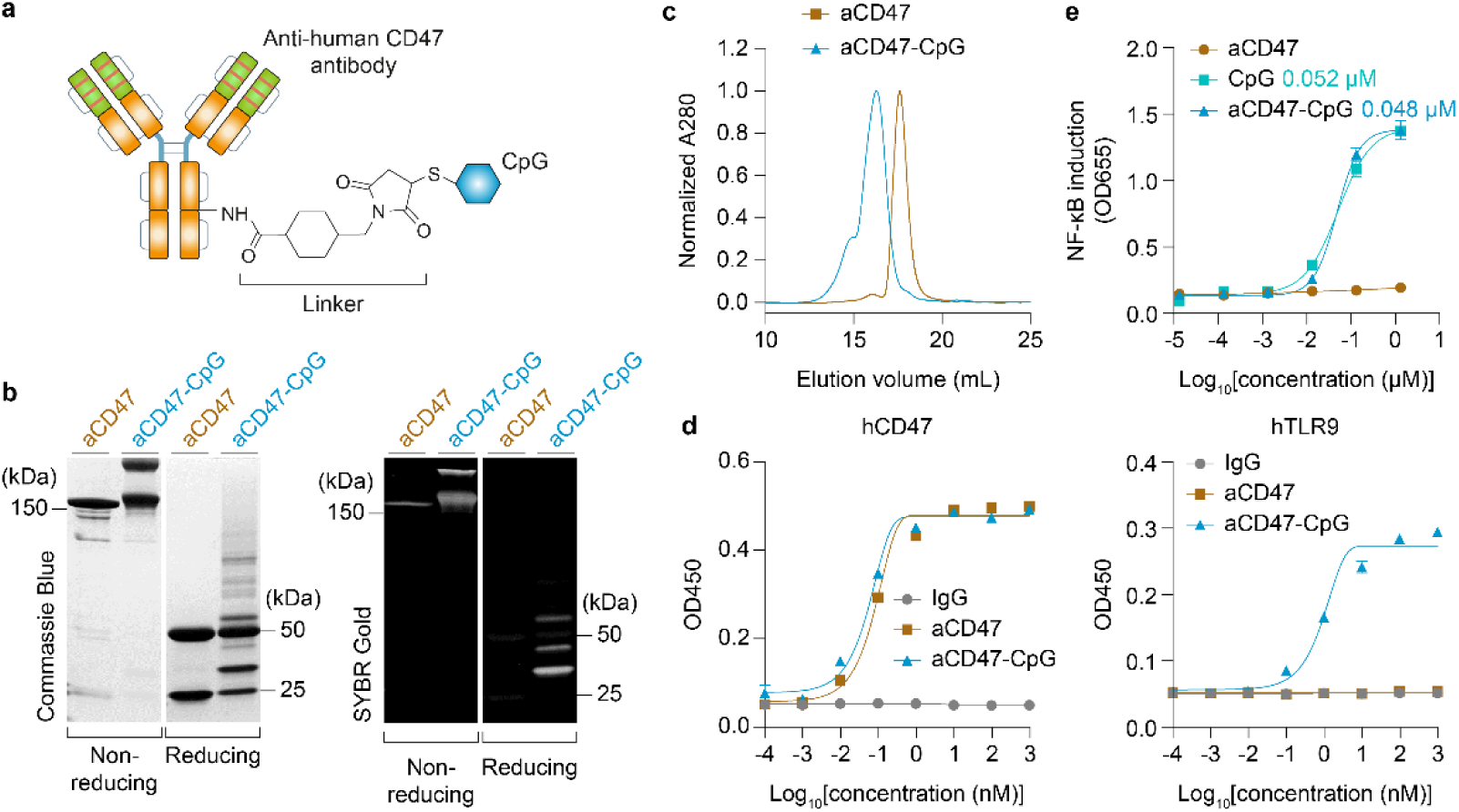
| Characterization of the human aCD47-CpG conjugate. **a**, Schematic illustration of the human aCD47-CpG engineered with human aCD47. **b**, SDS-PAGE analysis showing the conjugation patterns of human aCD47-CpG under non-reducing and reducing conditions. Left and right panels show Coomassie Blue and SYBR Gold staining for the visualization of aCD47 and CpG, respectively. **c**, Size exclusion chromatography analysis. **d**, Binding curves of aCD47-CpG to recombinant human CD47 (hCD47) and human TLR9 (hTLR9). **e**, NF-κB activity induced by aCD47-CpG.

**Extended Data Fig. 2.**
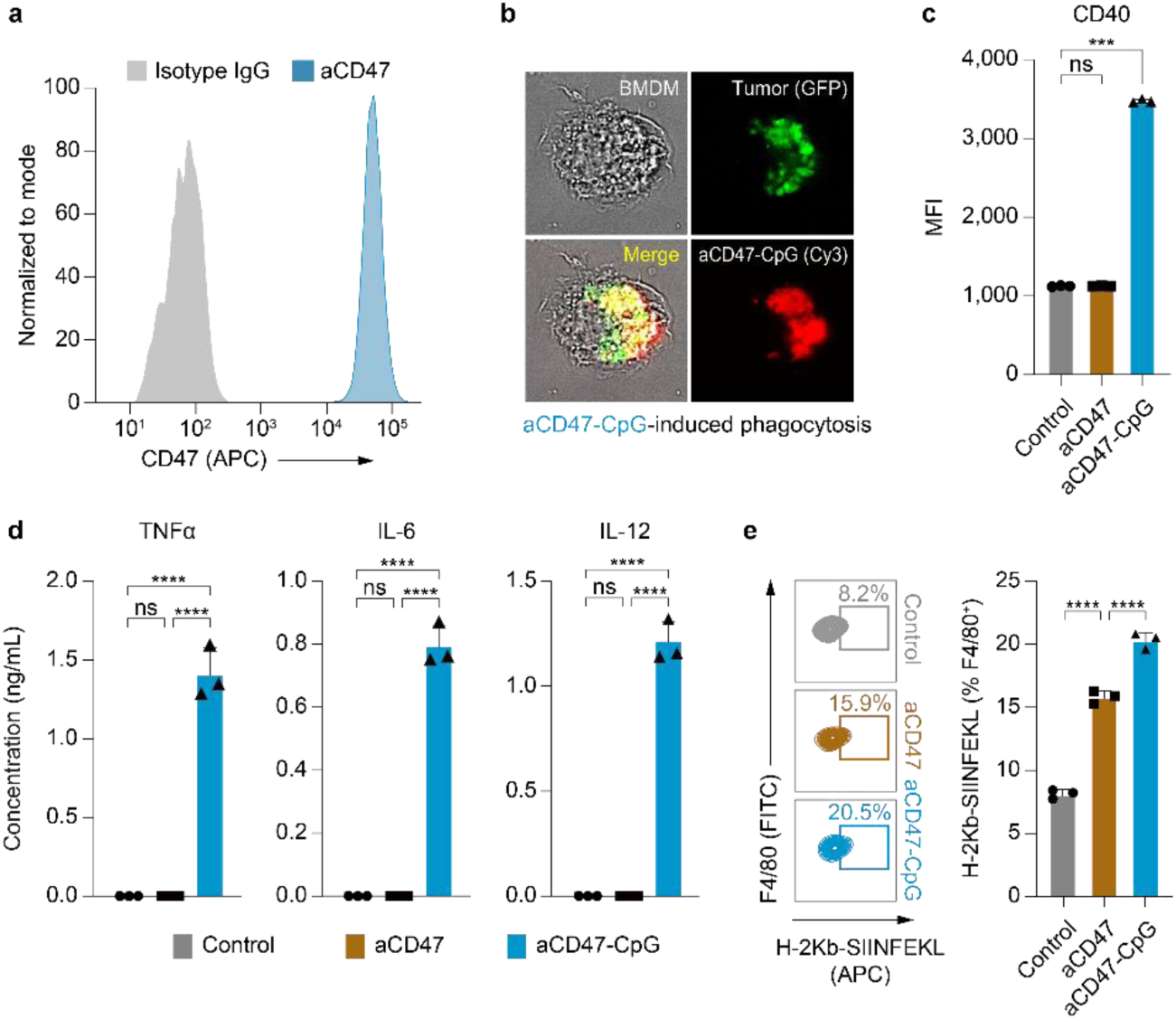
| Human aCD47-CpG promotes anti-tumor immune response of macrophages against Raji human lymphoma cells. **a**, Flow cytometry for expression of CD47 on human Raji lymphoma cells. Raji cells were stained with APC-labeled isotype IgG antibody (gray) or aCD47 (blue). **b**–**d,** BMDMs were co-cultured with Raji human lymphoma cells expressing GFP at a 2:1 ratio in the presence of aCD47 or aCD47-CpG. **b**, Representative fluorescence microscopy images showing aCD47-CpG-induced phagocytosis. BMDMs were co-cultured with Raji-GFP cells in the presence of Cy3-labeled aCD47-CpG. **c**, Expression of co-stimulatory molecule CD40 from BMDMs co-cultured with Raji cells after treatment with aCD47 or aCD47-CpG. **d**, Pro-inflammatory cytokines released from BMDMs by aCD47-CpG stimulation. The levels of TNFα, IL-6, or IL-12 secreted to cell culture supernatants were measured by ELISA after 24 h of incubation with aCD47 or aCD47-CpG. **e**, Cross-presentation of chicken ovalbumin-derived SIINFEKL peptide loaded onto H-2Kb (MHC-I) of macrophages. BMDMs were co-cultured with Raji cells expressing ovalbumin (Raji-OVA) in the presence of aCD47 or aCD47-CpG overnight at 37°C. Antigen cross-presentation was determined using flow cytometry by detecting MHC-I-bound SIINFEKL peptide on F4/80^+^ macrophages. Data shown are mean ± s.d. of three experimental replicates. ns: non-significant, \*\*\**p* < 0.001, \*\*\*\**p* < 0.0001 (one-way ANOVA with Tukey’s multiple comparisons test).

**Extended Data Fig. 3.**
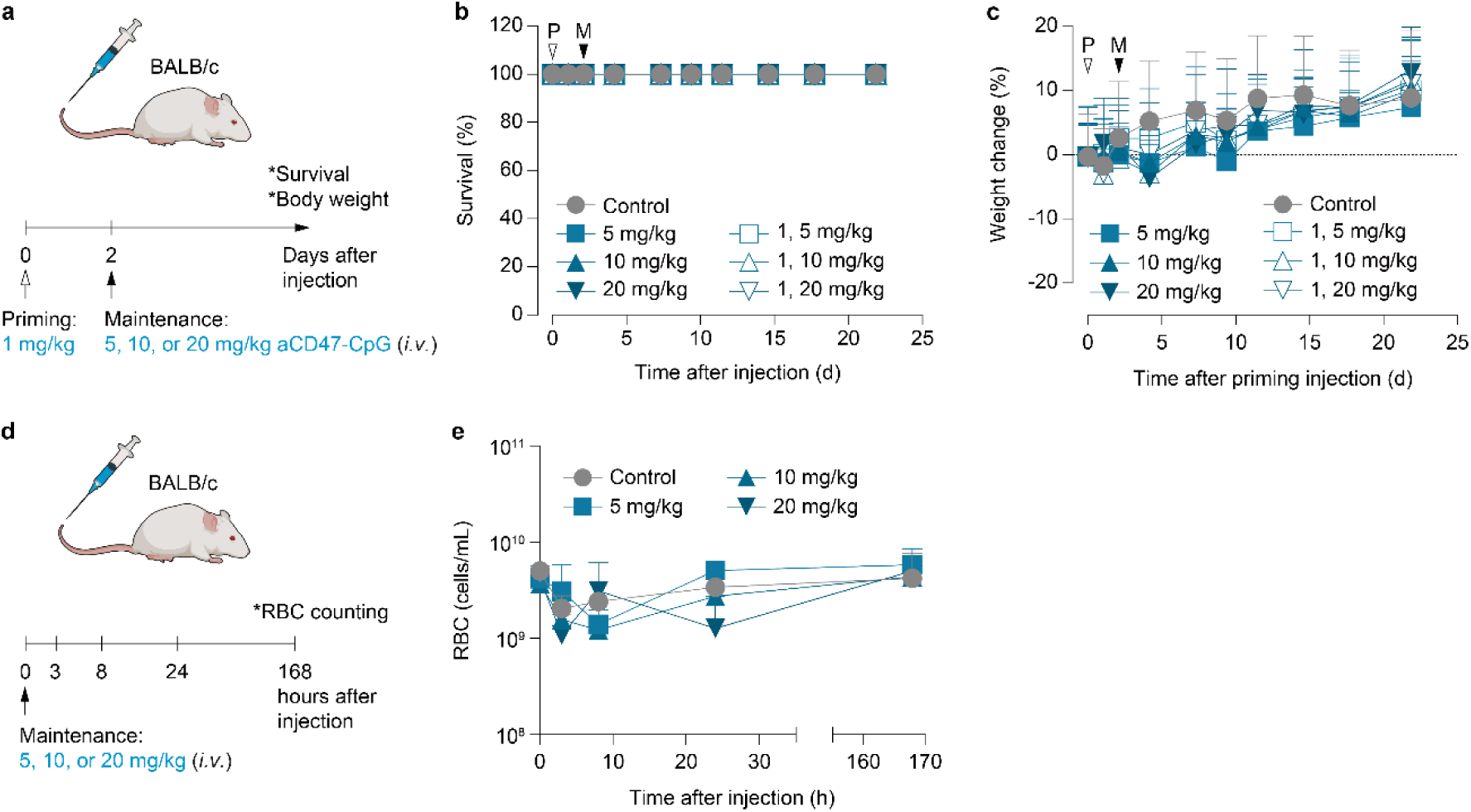
| Systemically administered mouse aCD47-CpG does not induce significant adverse effects. **a**, Schematic diagram of aCD47-CpG treatment in wild-type BALB/c mice for safety assessments. Animals were administered intravenously (*i.v.*) with a maintenance dose (M, black arrow) of aCD47-CpG at three escalating doses, with or without an *i.v.* priming dose (P, white arrow) at 1 mg/kg (*n* = 5 per group). **b**,**c**, Animals were monitored for survival (**b**) and body weight changes (**c**). **d**, Schematic diagram for hematotoxicity assessment. **e**, Red blood cell (RBC) counting using blood samples collected at the indicated time points from healthy BALB/c mice treated *i.v.* with aCD47-CpG at three escalating doses (*n* = 5 per group).

**Extended Data Fig. 4.**
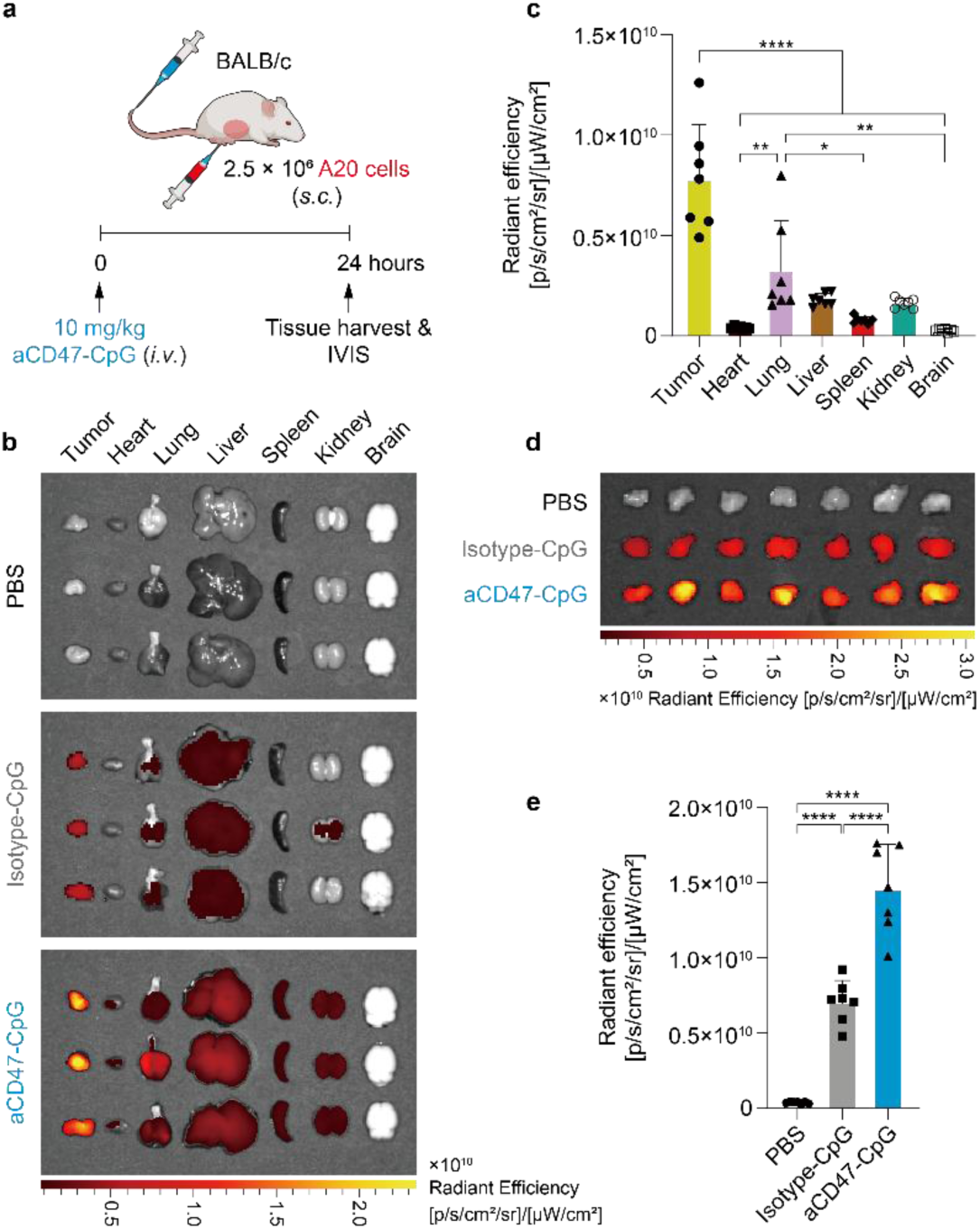
| Systemically administered mouse aCD47-CpG accumulates in tumors *in vivo*. **a**, Schematic illustration of the experimental schedule for the biodistribution study. BALB/c mice bearing A20 lymphomas (∼100–200 mm³) were intravenously (*i.v.*) administered with PBS, Cy5-labeled Isotype-CpG (isotype-CpG), or Cy5-labeled aCD47-CpG at a dose of 10 mg/kg. Tumors and major organs were harvested at 24-h post-injection for *ex vivo* imaging and analysis. **b**, Representative *ex vivo* fluorescence images of tumors and major organs (heart, lung, liver, spleen, kidney, and brain) harvested from mice in the PBS, isotype-CpG, and aCD47-CpG groups (*n* = 3 representative mice per group). **c**, Quantitative analysis of the biodistribution profile within the aCD47-CpG treated group (*n* = 7). The graph displays the fluorescence intensity of the tumor and indicated major organs. **d**, *Ex vivo* fluorescence images of all dissected tumors from the PBS, isotype-CpG, and aCD47-CpG groups (*n* = 7 per group). **e**, Quantification of aCD47-CpG accumulation in tumors compared to controls. Fluorescence signals were quantified as average radiant efficiency. Data are presented as mean ± s.d. \**p* < 0.05, \*\**p* < 0.01, \*\*\*\**p* < 0.0001 (one-way ANOVA with Tukey’s multiple comparisons test).

**Extended Data Fig. 5.**
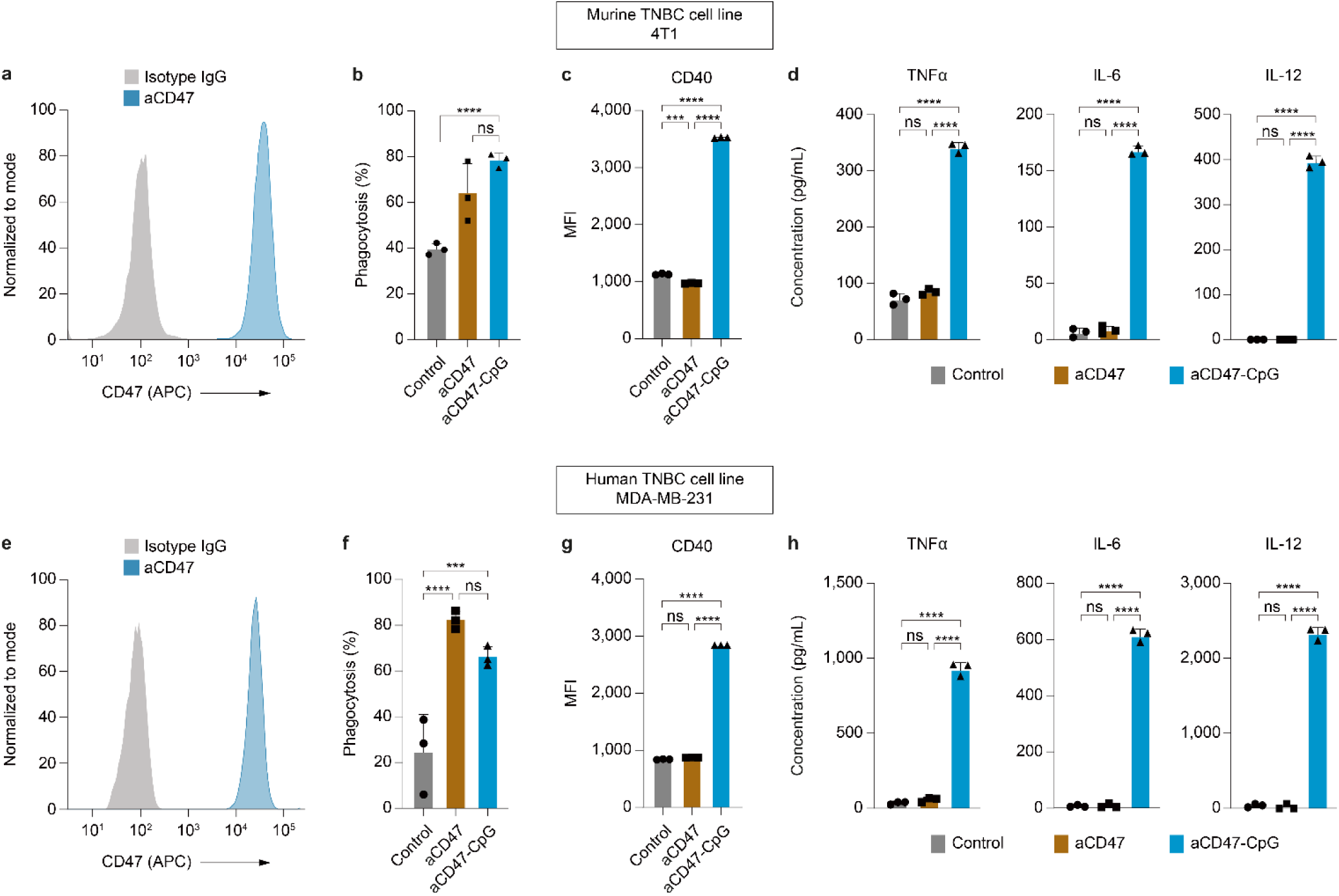
| aCD47-CpG promotes phagocytosis and anti-tumor immune response of macrophages against mouse and human TNBC cells. **a,e**, Flow cytometry for expression of CD47 on 4T1 (**a**) or MDA-MB-231 cells (**e**). Cells were stained with APC-labeled isotype IgG antibody (gray) or aCD47 (blue). **b**–**d**,**f**–**h**, BMDMs were co-cultured with 4T1 mouse (**b**–**d**) or MDA-MB-231 human breast cancer cells (**f**–**h**) at a 2:1 ratio in the presence of aCD47 or aCD47-CpG. **b**,**f**, Phagocytosis of luciferase-expressing 4T1 cells (**b**) or MDA-MB-231 cells (**f**) in the presence of aCD47 or aCD47-CpG overnight at 37°C determined by luminescence-based long-term macrophage killing (LB-LTMK) assay. **c**,**g**, Expression of co-stimulatory molecule CD40 from BMDM co-cultured with 4T1 cells (**c**) or MDA-MB-231 cells (**g**) after treatment with aCD47 or aCD47-CpG. **d**,**h**, Pro-inflammatory cytokines released from BMDM by aCD47-CpG stimulation. The levels of TNFα, IL-6, or IL-12 secreted to cell culture supernatants were measured by ELISA after 24 h of incubation with aCD47 or aCD47-CpG. Data shown are mean ± s.d. of three experimental replicates. ns: non-significant, \*\*\**p* < 0.001, \*\*\*\**p* < 0.0001 (one-way ANOVA with Tukey’s multiple comparisons test).

