## Supplementary Information for "An immune-stimulating antibody conjugate spatiotemporally targeting CD47 and TLR9 elicits macrophage-dependent tumor clearance and durable anti-cancer adaptive immunity"

This PDF file includes:

- **Supplementary Fig. 1** | Raw-Blue cells exhibit a moderate yet significant killing effect against A20 murine lymphoma cells overexpressing CD47, as demonstrated in luminescence-based assay.
- **Supplementary Fig. 2** | aCD47 and aCD47-CpG do not directly affect the luciferase activity of A20 cells.
- **Supplementary Fig. 3** | Systemic and intratumoral biodistribution of aCD47-CpG.
- **Supplementary Fig. 4** | Tumor growth of a localized TNBC syngeneic mouse model in each treatment group.
- **Supplementary Table 1** | $K_D$ values measured for mouse CD47 across aCD47-CpG conjugates with varying DAR (0–4).

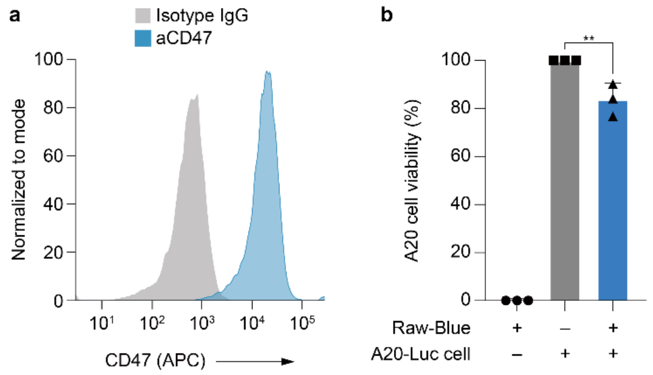


**Supplementary Fig. 1 | Raw-Blue cells exhibit a moderate yet significant killing effect against A20 murine lymphoma cells overexpressing CD47, as demonstrated in luminescence-based assay. a**, Flow cytometry analysis of the expression of CD47 on A20 murine lymphoma cell. A20 cells were stained with APC-labeled isotype IgG antibody (gray) or anti-CD47 antibody (blue). **b**, Luminescence-based long-term macrophage killing (LB-LTMK) assay using Raw-Blue cells and luciferase-expressing A20 (A20-Luc) cells. A20-Luc cells ($3 \times 10^4$) were co-cultured with Raw-Blue cells ($6 \times 10^4$) for 24 h at 37°C, followed by *in vitro* luciferase assay, where cells were lysed and then luminescence signal was detected using luciferase assay substrate. **$p < 0.01$ (one-way ANOVA with Tukey's multiple comparisons test).

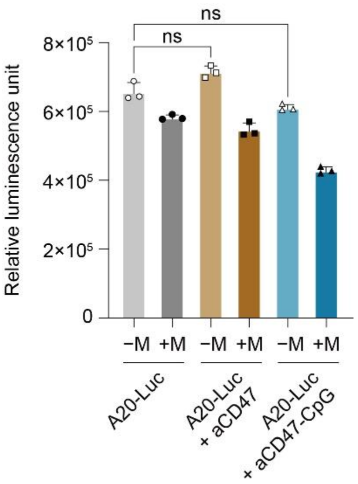


**Supplementary Fig. 2 | aCD47 and aCD47-CpG do not directly affect the luciferase activity of A20 cells.** Luciferase activity was measured in A20-Luc cells cultured either alone (−M) or co-incubated with macrophages (+M), following treatment with PBS, aCD47, or aCD47-CpG. The data confirm that the treatments do not intrinsically alter the luminescence signal in the absence of macrophages. ns: non-significant (one-way ANOVA with Tukey’s multiple comparisons test).

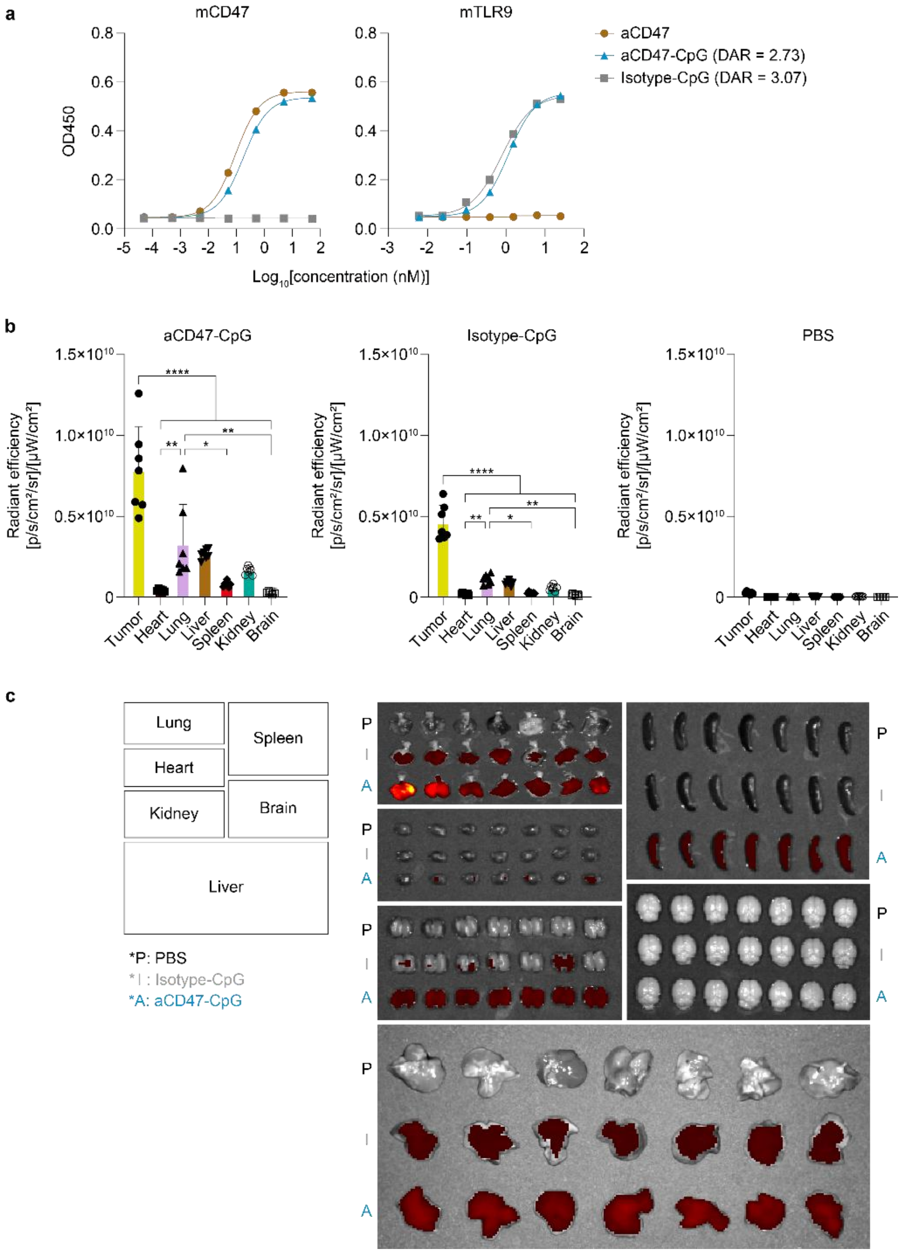


**Supplementary Fig. 3 | Systemic and intratumoral biodistribution of aCD47-CpG. a**, Binding curves of aCD47-CpG and isotype-CpG control to recombinant mouse CD47 (mCD47, left) and TLR9 (mTLR9, right). **b**, Quantitative analysis of *ex vivo* fluorescence intensity in tumors and major organs (heart, lung, liver, spleen, kidney, and brain) harvested 24 h post-injection. Mice were intravenously injected with PBS, Cy5-labeled isotype-CpG (10 mg/kg), or Cy5-labeled aCD47-CpG (10 mg/kg). Fluorescence signals were quantified as average radiant efficiency. **c**, *Ex vivo* fluorescence images of individual organs from all mice ($n$ = 7 per group). Data are presented as mean ± s.d. $^*p < 0.05$, $^{**}p < 0.01$, $^{****}p < 0.0001$ (one-way ANOVA with Tukey's multiple comparisons test).

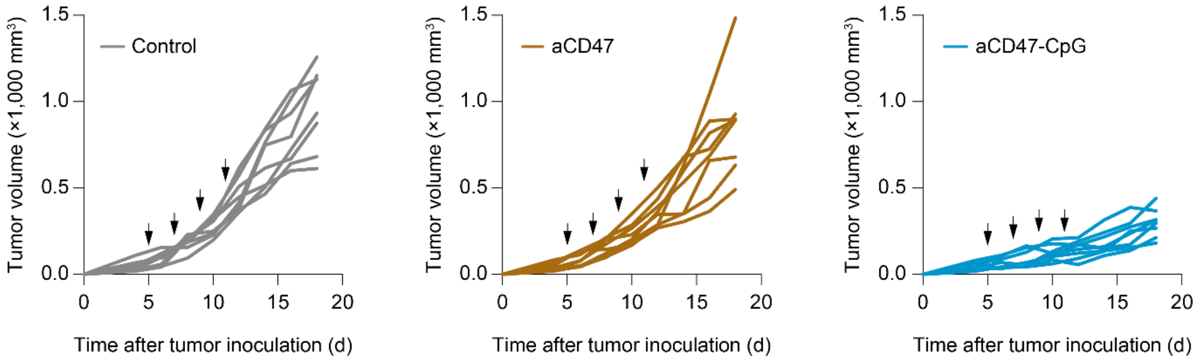


**Supplementary Fig. 4 | Tumor growth of a localized TNBC syngeneic mouse model in each treatment group.** Animals were treated intravenously with PBS, aCD47 (10 mg/kg), or aCD47-CpG (10 mg/kg) every third day up to four doses (black arrows). Tumor volume measured using a caliper and presented for each individual mouse.

| aCD47-CpG | $K_D$ value (M, × $10^{-10}$) |
|---|---|
| DAR0 | 1.464 |
| DAR1 | 1.678 |
| DAR2 | 2.270 |
| DAR3 | 3.185 |
| DAR4 | 3.274 |

**Supplementary Table 1. $K_D$ values measured for mouse CD47 across aCD47-CpG conjugates with varying DAR (0–4).** The $K_D$ values were calculated from ELISA investigating binding properties of aCD47-CpG with each DAR to mouse CD47.
